# Phylogenomic terraces: presence and implication in species tree estimation from gene trees

**DOI:** 10.1101/2020.04.19.048843

**Authors:** Ishrat Tanzila Farah, Md Muktadirul Islam, Kazi Tasnim Zinat, Atif Hasan Rahman, Md Shamsuzzoha Bayzid

## Abstract

Species tree estimation from multi-locus dataset is extremely challenging, especially in the presence of gene tree heterogeneity across the genome due to incomplete lineage sorting (ILS). *Summary methods* have been developed which estimate gene trees and then combine the gene trees to estimate a species tree by optimizing various optimization scores. In this study, we have formalized the concept of “phylogenomic terraces” in the species tree space, where multiple species trees with distinct topologies may have exactly the same optimization score (*quartet score, extra lineage score*, etc.) with respect to a collection of gene trees. We investigated the presence and implication of terraces in species tree estimation from multi-locus data by taking ILS into account. We analyzed two of the most popular ILS-aware optimization criteria: *maximize quartet consistency* (MQC) and *minimize deep coalescence* (MDC). Methods based on MQC are provably statistically consistent, whereas MDC is not a consistent criterion for species tree estimation. Our experiments, on a collection of dataset simulated under ILS, indicate that MDC-based methods may achieve competitive or identical quartet consistency score as MQC but could be significantly worse than MQC in terms of tree accuracy – demonstrating the presence and affect of phylogenomic terraces. This is the first known study that formalizes the concept of phylogenomic terraces in the context of species tree estimation from multi-locus data, and reports the presence and implications of terraces in species tree estimation under ILS.

## Introduction

A *species tree* can be defined as the pattern of branching of species lineages via the process of speciation, while a *gene tree* represents the evolution of a particular “gene” within a group of species. Biological processes can result in different loci having different evolutionary histories, and therefore species tree estimation involves the estimation of trees and alignments on many different genes, so that the species tree can be based upon many different parts of the genome. While many processes can result in discord between gene trees and species trees, incomplete lineage sorting (ILS) is considered to be a dominant cause for gene tree heterogeneity, which is best understood under the coalescent model [1–8]. ILS or deep coalescence refers to the case in which two lineages fail to coalesce at their speciation point. Under the coalescent model, deep coalescence can be a source of discordance, because the common ancestry of gene copies at a single locus can extend deeper than speciation events.

In the presence of gene tree heterogeneity, standard methods for estimating species trees, such as concatenation (which combines sequence alignments from different loci into a single “supermatrix”, and then computes a tree on the supermatrix) can be statistically inconsistent [9, 10], and produce incorrect trees with high support [11]. Therefore, methods for estimating species trees that can explicitly take ILS into account have been developed, and some of them are provably statistically consistent meaning that they will converge in probability to the true species tree given sufficiently large numbers of genes and sites per gene. Many of these methods operate by combining estimated gene trees, and so are called “summary methods”. Examples of statistically consistent coalescent based summary methods include MP-EST [12], ASTRAL [13, 14], BUCKy [15], GLASS [16], STEM [17], SVDquartets [18], STEAC [19], NJst [20], ASTRID [21], STELAR [22]. Other statistically consistent species-tree estimation methods include BEST [23] and *BEAST [24], which co-estimate gene trees and species trees from input sequence alignments. These methods can produce substantially more accurate trees that other methods; however, these methods are extremely computationally intensive and do not scale to large numbers of genes [25–27], so that only summary methods are feasible for use on genomescale datasets.

Maximize quartet consistency (MQC) is one of the leading optimization criteria for estimating statistically consistent species trees from gene trees in the presence of ILS [13, 28–30]. MQC seeks for a species tree that is consistent with the largest number of quartets induced by the set of gene trees. ASTRAL [13, 14], which is one of the most accurate and popular coalescent based summary methods, solves this optimization problem. Since MQC is an NP-hard problem, ASTRAL has two versions: the exact version is guaranteed to return the globally optimal solution but runs in exponential time, and the heuristic version returns the optimal tree for a constrained search space and runs in polynomial time.

Another optimization problem that takes ILS into account is minimize deep coalescence (MDC), which was first introduced by Maddison in his seminal paper [31] and was further investigated by [32, 33]. In the presence of gene tree discordance due to ILS, MDC criterion seeks for species trees that minimize the number of “deep” coalescence events required for a given collection of gene trees [32, 34, 35]. Given a collection of gene trees *𝒢* = {*gt*_1_, *gt*_2_, *…, gt*_*k*_}, MDC seeks a species tree *T* such that the total number of extra lineages (ELs) with respect to *𝒢*, denoted by *XL*(*T, 𝒢*) = Σ*i*(*XL*(*T, gt*_*i*_)), is minimized. Here, *XL*(*T, gt*_*i*_) denotes the number of extra lineages of a species tree *T* with respect to a gene tree *gt*_*i*_. This is defined by embedding the gene tree *gt*_*i*_ into the species tree *T*, and then counting the number of lineages on each edge of the species tree. The number of extra lineages on an edge is one less than the total number of lineages on that edge [36]. Thus, it is a natural candidate for species tree inference, under the parsimony setting, when discordance among gene trees is caused by incomplete lineage sorting. Several exact algorithms and heuristics for implementing this criterion have been developed [32,37]. Phylonet [38] is a popular tool for estimating species trees under MDC, which has both exact and heuristic versions similar to ASTRAL. Although it has been shown to be statistically inconsistent under multi-species coalescent model [39], this is not agnostic to the gene tree heterogeneity as it takes into account the specific nature of the way in which incomplete lineage sorting occurs. Simulation studies have suggested a high degree of accuracy of species tree estimates obtained by this criterion [32, 40]. Yet, to our knowledge, the accuracy of species trees estimated under MDC has not been explored in comparison to highly accurate ILS-aware and statistically consistent summary methods like ASTRAL. Although statistically consistent methods like ASTRAL is expected to perform better than statistically inconsistent methods like Phylonet, it is important to evaluate the relative performance under various realistic model conditions as even the best coalescent-based summary methods are sensitive to gene tree estimation error and have not been reliably more accurate than concatenation [25, 41]. There have been a few studies [13, 25, 42, 43] which present comparisons among concatenation and various summary methods, including ASTRAL, MP-EST, BUCKy, NJst, and SVDquartets. Huang *et al.* [44] evaluated STEM and Phylonet in the presence of mutational and coalescent variance. However, to the best of our knowledge, there is no study evaluating MQC and MDC.

This study shows that MQC is in general a better optimization criterion compared to MDC. However, MDC achieves competitive results on some of the model conditions that we analyzed in this study. Interestingly, this study reveals that the search under MDC criterion may result into trees that have competitive *quartet score (QS)* (number of quartets induced by the gene trees that a tree is consistent with) compared to the trees estimated by ASTRAL. However, MDC trees are generally significantly worse than ASTRAL in terms of tree accuracy. This could be due to the presence of “islands” of topologically different trees with equally good optimization score. Therefore, we have formalized the concept of “phylogenomic terraces”, where potentially large numbers of distinct species trees may have exactly the same optimization score with respect to a set of input gene trees. In particular, this study entails the following contributions.

- We have introduced and formalized the concept of phylogenomic terraces in the context of constructing species trees from a collection of gene trees.
- We proved important structural properties showing that phylogenomic terraces are obvious especially when we have a large number of taxa. We proved relevant theoretical results for “MDC-terrace” and “MQC-terrace”.
- We investigated MDC (Phylonet-MDC) and MQC (ASTRAL) criteria in terms of tree accuracy under different model conditions with varying numbers of genes, amounts of ILS and gene sequence lengths. We also investigated their performance on real biological dataset.
- We systematically analyzed, through simulation studies, the impact of terraces in the search for optimal species trees under MDC and MQC criteria.

## Terraces in Phylogenomic Analyses

Estimating species trees from multiple markers, in the presence of gene tree heterogeneity, starts with estimating gene trees from individual gene sequence alignments and then summarizes them to get a coherent species tree. Summarizing is typically done by optimizing various optimization criteria, e.g minimizing deep coalescence [31, 32, 45], maximizing pseudo-likelihood [12], maximizing quartet consistency [13, 28] etc. Since the tree space grows exponentially as the number of taxa increases, navigating through the tree space to find the optimal tree is challenging. Sanderson *et al.* showed that, when phylogenetic trees are estimated from sequence alignments using maximum likelihood (ML), multiple distinct trees can have exactly the same likelihood score – a condition which was referred to as terraces [46] and was further investigated in subsequent studies [47–49]. A similar concept is “islands” of trees under ML and MP (maximum parsimony) criteria for estimating trees from sequence data [50, 51]. Phylogenetic tree islands have been explored in the context of single-locus data. Sanderson *et al.* [46] considered multi-locus data but they analyzed the supermatrix resulting from concatenating the gene alignments, and characterized the terraces in terms of the ML score with respect to the combined alignments. Similar phenomenon can arise in phylogenomic analyses where species trees are estimated from a set of gene trees under various optimization criteria. However, terraces in this context has never been explored before.

We formalize the concept of terraces for phylogenomic analyses under various optimization criteria. For an optimization criterion *𝒞*, we denote by *𝒞*-*terrace* a set of trees with distinct tree topologies but with exactly the same *𝒞*-*score*.

**Definition 1.** *For a set 𝒢 of gene trees and an optimization criterion 𝒞, the 𝒞-terrace T*_*𝒢,𝒞*_(*s*) *represents a set of species trees having exactly the same 𝒞-score s with respect to the input set 𝒢 of gene trees.*

In this paper, we particularly investigate MDC- and Quartet-terrace. We prove theoretical results showing the possibility of the existence of MDC- and quartet-terraces, especially when the number of taxa is high.

**Theorem 2.** *For a set 𝒢 of k gene trees on n taxa, the species tree space will have at least one MDC-terrace if* 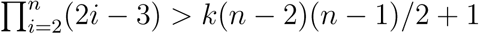.

*Proof.* For a gene tree *gt* and a species tree *ST* on *n* taxa, the number of extra lineages increases as the number of deep coalescence increases, that means the lineages do not coalesce at their common ancestors and go further deeper in time. Therefore, the number of extra lineages is maximized when the gene lineages do not coalesce until they reach the root of the phylogeny, resulting into extra lineages on all the internal branches. Assuming that gene lineages do not coalesce with each other until they reach the root node, the number of extra lineages will be maximized for a pectinate tree (also known as caterpillar tree) since all the gene lineages (except for one) will go through all the internal branches “above” its most immediate ancestor. See Figure 1 for an example which shows how a pectinate species tree results into more extra lineages than relatively more balanced trees. Therefore, the internal branch incident on the root of the phylogeny will contain *n* – 1 lineages and hence *n* − 2 extra lineages. The more recent internal branches (from ancient to recent) will contain *n* − 3, *n* − 4, …, 1 extra lineages, respectively. Thus, maximum number of extra lineage that may occur for a gene tree *gt* and a species tree *ST* is

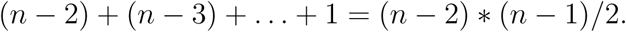

For a set *𝒢* of *k* gene trees, maximum number of extra lineages with respect to a species tree *ST* will be *k*(*n* − 2)(*n* − 1)*/*2. The number of rooted species trees in the tree space with *n* taxa (*n* ≥ 2) is

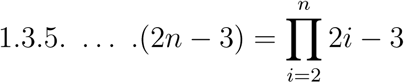

[52, 53]. The MDC-scores of these trees in the species tree space will be within the range 0 ∼ *k*(*n* − 2)(*n* − 1)*/*2. That means there are k(n-2)(n-1)/2 + 1 possible distinct MDC-scores for 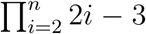 trees in the tree space with respect to *𝒢*. Therefore, using pigeonhole principle, there will be at least one terrace *𝒯*_*G*_ with more than one trees having identical MDC-score provided that 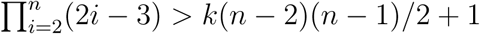.

**Figure 1:**
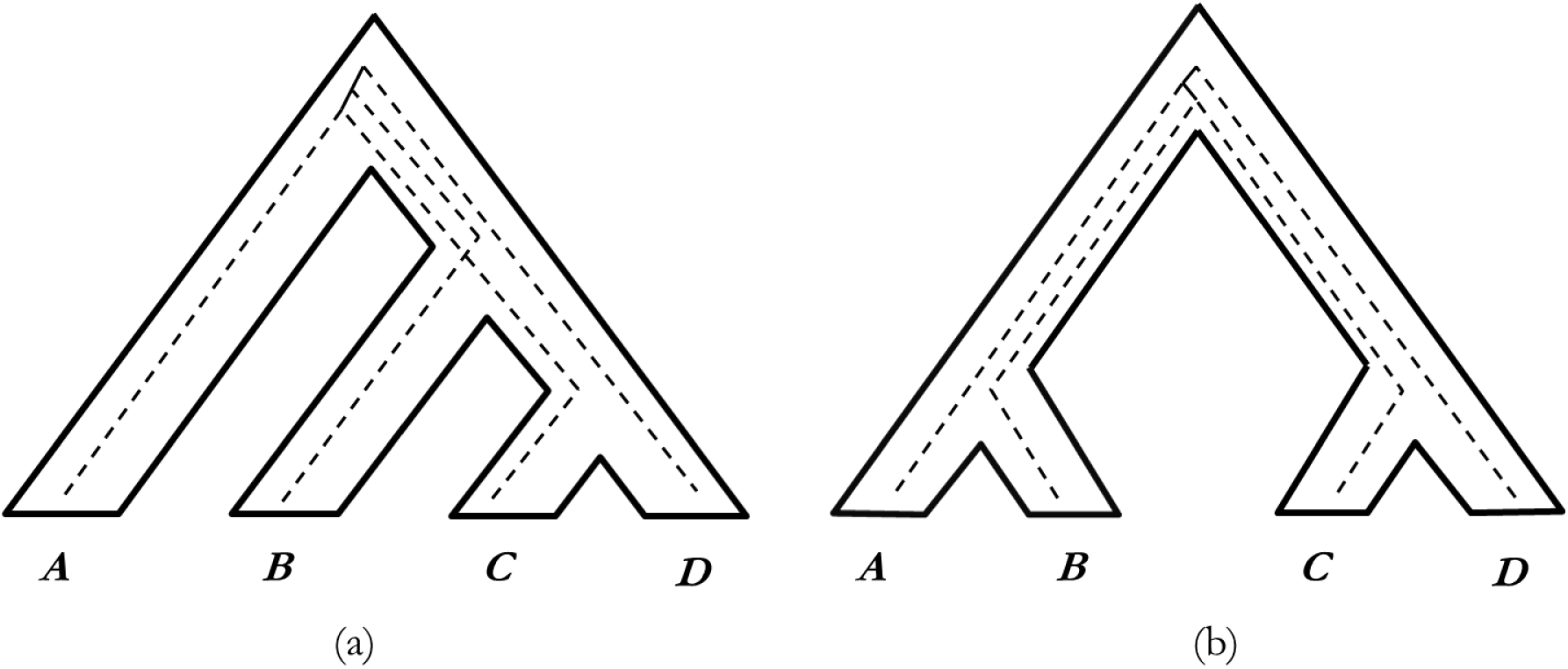
Extra lineages inside a pectinate and a balanced symmetric tree for the cases when gene lineages do not coalesce with each other on the internal branches until they reach the root of the tree. Species boundaries are shown in solid lines while gene lineages are represented by dashed lines. (a) Reconciliation of *gt* = (((*A, C*), *B*), *D*) with a pectinate species tree *ST* = (((*A, B*), *C*), *D*) which results into three extra lineages. (b) Reconciliation of *gt* = (((*B, C*), *A*), *D*) with a balanced species tree *ST* = ((*A, B*), (*C, D*)) which results into two extra lineages. In both cases, the gene lineages do not coalesce with each other until they go backwards to the root node.

Table 1 shows an example demonstrating the MDC-terraces in the tree space for four taxa with respect to a set *𝒢* of four rooted gene trees. There are 15 possible rooted species tree topologies with four taxa and we examined the MDC scores (extra lineage score) of all of them with respect to *𝒢*. There are eight terraces: *𝒯*_*𝒢,ℳ𝒢𝒞*_(4), *𝒯*_*𝒢,ℳ𝒢𝒞*_(5), *𝒯*_*𝒢,ℳ𝒢𝒞*_(6), *𝒯*_*𝒢,ℳ𝒢𝒞*_(7), *𝒯*_*𝒢,ℳ𝒢𝒞*_(8), *𝒯*_*𝒢,ℳ𝒢𝒞*_(9), *𝒯*_*𝒢,ℳ𝒢𝒞*_(10), and *𝒯*_*𝒢,ℳ𝒢𝒞*_(11) for this given set of gene trees with MDC scores of 4, 5, 6, 7, 8, 9, 10 and 11. Here, two candidate species trees (((*B, D*), *C*), *A*) and (((*C, D*), *B*), *A*) belong to *𝒯*_*𝒢,ℳ𝒢𝒞*_(4) and are optimal (most parsimonious) under the MDC score. This multiplicity of optimal trees under a particular optimization criteria introduces ambiguity to tree search algorithms. Since the number of possible tree topologies increases exponentially as the number of taxa increases whereas the MDC scores grows only at a polynomial rate, it is expected from Theorem 2 that the number of MDC terraces will grow as we increase the number of taxa.

**Table 1:**
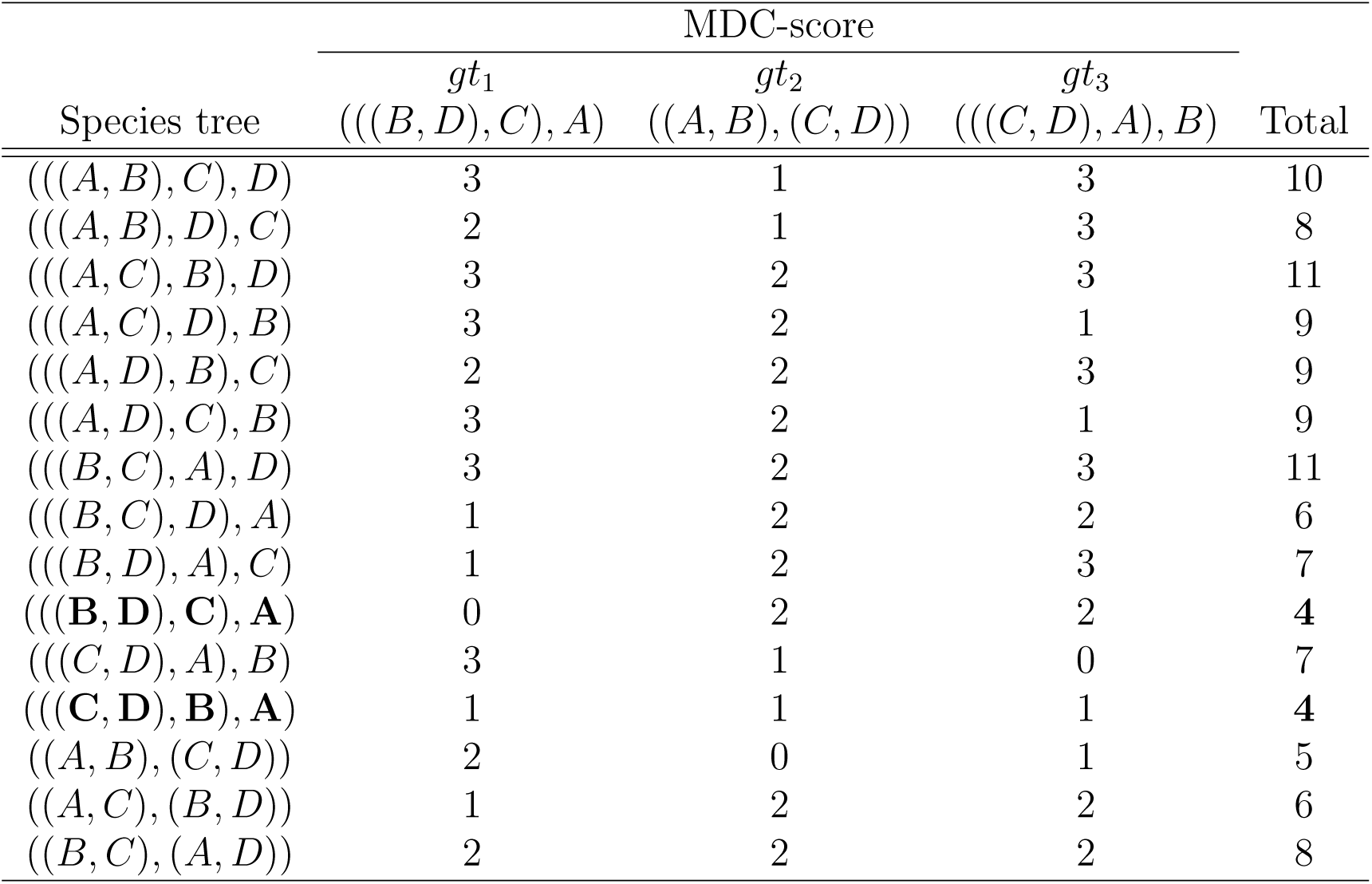
MDC-terraces in the tree space of four taxa with respect to a set *𝒢* of rooted and binary gene trees. *𝒢* contains four gene trees, where two of the genes have the gene tree topology of *gt*_1_ = (((*B, D*), *C*), *A*), one gene tree has the topology of *gt*_2_ = ((*A, B*), (*C, D*)), and one gene has the topology of *gt*_3_ = (((*C, D*), *A*), *B*). Two species trees (((*B, D*), *C*), *A*) and (((*C, D*), *B*), *A*) are both optimal under MDC with 4 extra lineages.

**Theorem 3.** *For a set 𝒢 of k gene trees on n taxa, the species tree space will have at least one quartet-terrace if* 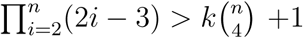.

*Proof.* There are 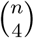 quartets in a tree with *n* taxa. Therefore, for a gene tree *gt* ∈ *𝒢* and a species tree *ST*, both on the same set of *n* taxa, *ST* can satisfy at most 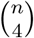 quartets (this is when *gt* and *ST* have identical topology). Hence, for a set *𝒢* of *k* gene trees, maximum number of consistent quartets with respect to a species tree *ST* is *k* 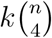. Following the same argument as described in Theorem 3, there will be at least one quartet-terrace *𝒯*_*Q*_ with more than one trees having identical quartet-score if 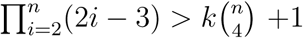. □

Note that the numbers of distinct MDC-score and quartet-score grow at a polynomial rate, whereas the number of unique species trees grows exponentially as we increase the number of taxa. This is also true for most of the other optimization criteria (e.g., minimize gene duplication and loss (MGDL) score [54–56]) that are being frequently used in phylogenomic analyses. Thus, search for an optimal species tree under a particular optimization criteria may result into a tree which is a member of a terrace with potentially large number of trees with different tree topologies. Therefore, we may have multiple trees having the optimal (or near-optimal) score, but with different “closeness” to the true tree. Our experimental results also support this as we will show that, in some cases, the trees estimated by Phylonet have competitive or identical quartet scores, but are not as accurate as ASTRAL.

## Experimental Studies

### Datasets

We used previously studied simulated and biological datasets to evaluate the performance of MQC and MDC. We studied four collections of simulated datasets: one based on a biological dataset (37-taxon mammalian dataset) that was generated in a prior study [57], and three other simulated datasets from [57, 58]. Table 2 presents a summary of these datasets, under various model conditions with varying numbers of taxa (10 to 37), ILS levels (reflected in the average topological distance between true gene trees and true species tree) and gene sequence lengths.

**Table 2:** Properties of the simulated datasets. Level of ILS is presented in terms of the average topological distance between true gene trees and true species tree.

| Dataset | ILS level | # genes | # sites | # Replicates | Ref. |
| --- | --- | --- | --- | --- | --- |
| 10-taxon | 40%, 84% | 200 | 100 | 20 | [60] |
| 11-taxon | 38%, 85% | 5 - 100 | 2000 | 20 | [58] |
| 15-taxon | 82% | 100 - 1000 | 100 - 1000 | 10 | [60] |
| 37-taxon | 18%, 32%, 54% | 25 - 800 | 250 - 1000 | 20 | [57] |

In the mammalian simulation, we explored the impact of varying numbers of gene, the impact of phylogenetic signal by varying the sequence length (250bp, 500bp, 1000bp and 1500bp) for the markers. In both cases, three levels of ILS are simulated by multiplying or dividing all internal branch lengths in the model species tree by two, and we also explore various numbers of genes. These datasets have been generated under a multistage simulation process: a species tree was estimated on a mammalian dataset [59] using MP-EST, gene trees were simulated down the species tree under the multi-species coalescent model and then gene sequence alignments were simulated down the gene trees under the GTRGAMMA model.

We used three other simulated datasets: 11-taxon dataset (generated by [58] and subsequently studied by [25, 27]), 10-taxon and 15-taxon datasets (generated and studied by [57]). 10- and 11-taxon datasets vary in the levels of ILS (low and high amount of ILS; see Table 2). 15-taxon datasets vary in sequence length with high amount of ILS. Thus, the simulated datasets provide a range of conditions in which we explore the performance of MQC and MDC and investigated the impact of phylogenomic terrace.

We used two biological datasets: the 37-taxon mammalian datasets studied by Song *et al.* [59] with 424 genes, and and the the amniota dataset from [61] containing 16 species and 248 genes.

### Methods

We compared ASTRAL-III [62] which is based on MQC problem with Phylonet-MDC [32, 38] which is based on MDC. We ran the exact versions of ASTRAL and Phylonet on datasets with 10 ∼ 15 taxa, and the heuristic versions for larger dataset (37-taxon).

### Measurements

We compared the quartet support scores, extra lineage scores (number of extra lineages required to reconcile the input set of gene trees with a species tree [31,32]) and topological accuracy of trees computed by ASTRAL and Phylonet. We used false negative (FN) rate to measure the topological error. All the trees estimated by ASTRAL and Phylonet in this study are binary and so False Positive (FP) rate and False Negative (FN) rate are identical. For the biological dataset, we compared the estimated species trees with the existing literature and biological beliefs. We performed Wilcoxon signed-rank test (with *α* = 0.05) to measure the statistical significance of the differences between two methods.

## Results and Discussion

### Results on biologically based simulated datasets (37-taxon mammalian)

#### Missing branch rate

Substantial differences were observed between ASTRAL and Phylonet-MDC on all the model conditions that we analyzed. Figure 2 (a) shows the average FN rates on three model conditions with varying amounts of ILS. Both ASTRAL and Phylonet incurred the highest amount of missing branch rate (around 5% and 20% respectively) for 0.5X model condition which has the highest amount of gene tree discordance. ASTRAL has the lowest amount of FN rate (2.5%) for 2X model condition, which is expected as it has the lowest level of ILS. However, Phylonet was better on 1X model condition than 2X model condition. Figure 2(b) shows the error rates on various model conditions with varying lengths of gene sequence alignments and hence varying amounts of gene tree estimation error. We also analyzed the true gene trees. The highest amount of errors were observed on the model condition with the shortest sequences (i.e., highest amount of gene tree estimation error). The difference between ASTRAL and Phylonet-MDC was substantial (6.32% vs. 21.47%). As we increase the sequence lengths and hence decrease the gene tree estimation errors, both methods produced more accurate trees, but the differences between ASTRAL and Phylonet-MDC were still very substantial. Additionally, Phynonet incurred higher FN rate on 1000bp (10.29%) than on 500bp (9.26%). Even on the true gene trees, Phylonet was much worse than ASTRAL. Figure 2(c) shows the FN rates for varying numbers of genes. ASTRAL, being a statistically consistent methods, showed improved accuracy as we increase the number of genes. However, similar trend was not observed for Phylonet as increasing genes from 100 to 800 did not improve the tree accuracy. For 50, 100, 400 and 800 genes, Phylonet achieved very similar missing branch rates (12.35% ∼ 12.65%). Overall observation is that ASTRAL is much better in terms of FN rate than Phylonet and the differences are statistically significant (*P <<* 0.05).

**Figure 2:**
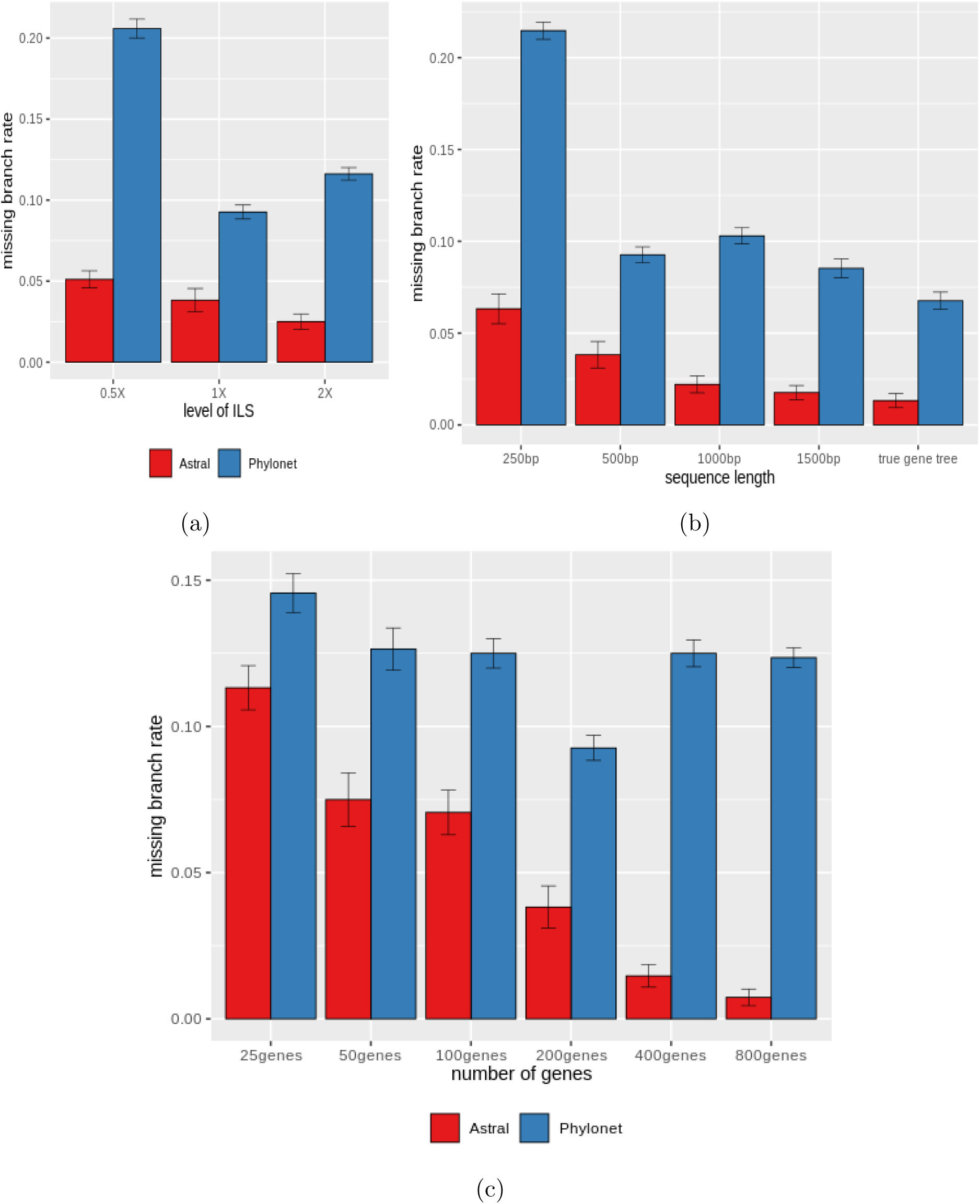
Comparison of ASTRAL and Phylonet on 37-taxon simulated mammalian dataset. We show the average FN rates with standard error bars over 20 replicates. (a) We fixed the sequence length to 500 bp and number of genes to 200, and varied the amounts of ILS. 2X model condition contains the lowest amount of ILS while 0.5X refers to the model condition with the highest amount of ILS. (b) We varied the amount of gene tree estimation error by varying the sequence lengths from 250 to 1000 bp, while keeping the ILS level (moderate) and the number of genes (200) fixed. (c) We fixed the sequence length to 500 bp and amount of ILS to 1X, and varied the numbers of genes from 25 ∼ 800).

#### Quartet Score

Figure 3 shows the comparison of the quartet scores of the species trees estimated by ASTRAL and Phylonet. We also show the quartet score of the true species tree (denoted by “true quartet score”). As expected, ASTRAL achieved higher quartet scores than Phylonet under all model conditions as ASTRAL estimates species trees by maximizing the quartet score. The difference with the true quartet score is less for ASTRAL than Phylonet. However, we observed that ASTRAL generally overestimates the quartet scores (compared to the true quartet score) except for a couple of model conditions (250 bp in Fig. 3(b) and 2X in Fig. 3(a)). For all the model conditions, Phylonet, in general, underestimated the quartet score since it does not take quartet consistency into account. However, Phylonet estimated trees are not as substantially different than the true trees with respect to quartet score as they are with regards to the topological accuracy. Interestingly, Phylonet achieved closer quartet scores (with respect to the true quartet scores) than ASTRAL (on 25- and 50-gene model conditions shown in Figure 3(c)). ASTRAL’s average quartet scores are higher than true quartet scores by 4308 and 4415 on 25-and 50-gene model conditions, whereas the average quartet scores of Phylonet are lower than the true quartet scores by only 122 and 516 for 25 and 50 genes, respectively. However, Phylonet is significantly worse than ASTRAL on these two model conditions (see Fig. 2 (c)) – suggesting that overestimation of quartet scores does not hurt the tree accuracy as much as underestimation. These results suggest that MDC criteria may achieve reasonably high QS, but the search under MDC may lead to incorrect trees by underestimating the amount of extra lineages. This indicates the presence of Quartet-terrace which we will discuss later in this section.

**Figure 3:**
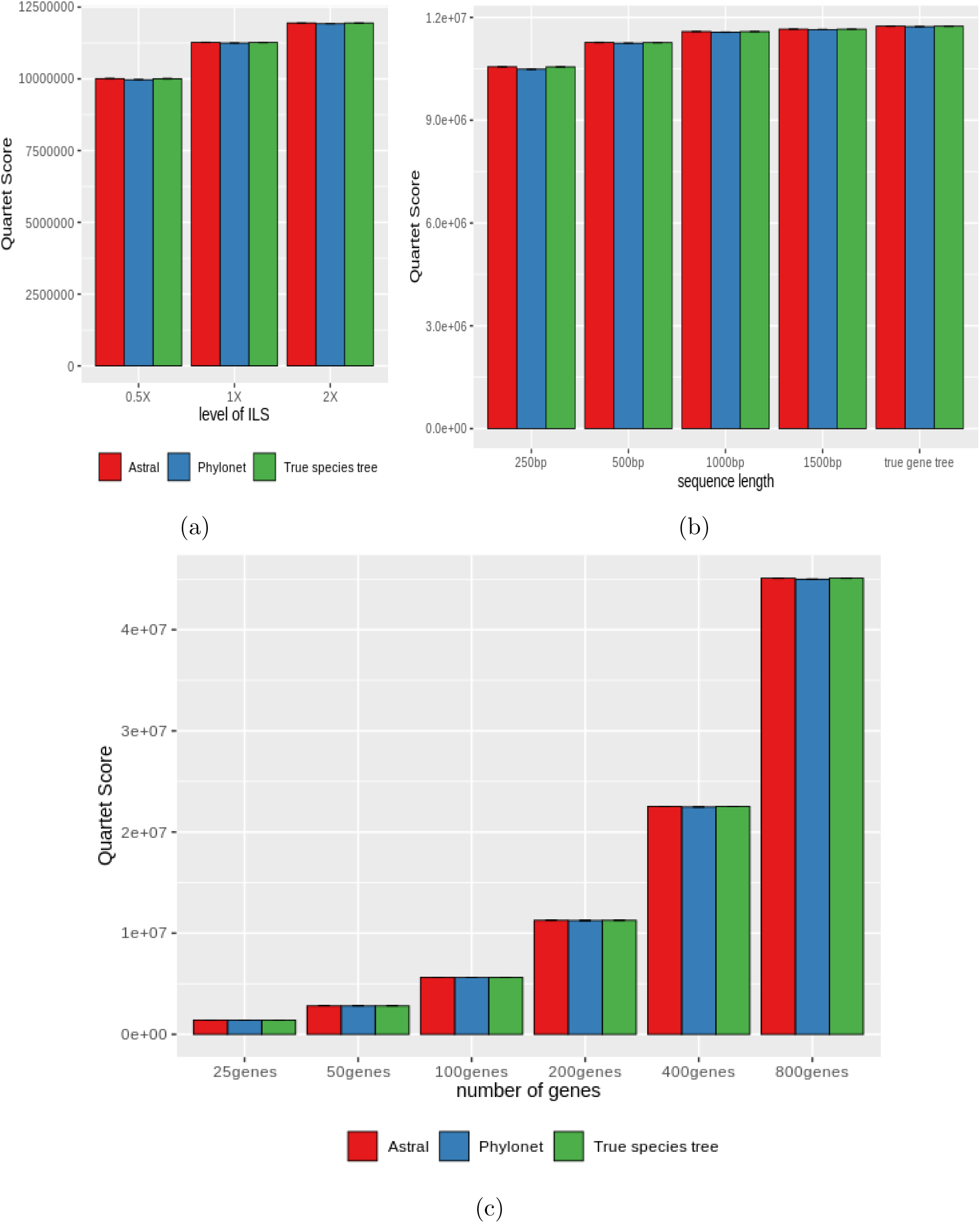
Quartet scores for ASTRAL, Phylonet and true species tree on biologically based simulated 37-taxon mammalian dataset. We show average quartet scores (number of quartets in the gene trees that are consistent with the species tree) with standard error bars over 20 replicates for various model conditions by controlling the levels of ILS, gene tree estimation error and numbers of genes.

#### Extra Lineage (EL) Score

We show the comparison between ASTRAL and Phylonet-MDC in terms of the number of extra lineages in Figure 4. We also show the EL scores of the true species trees (which we refer to as”true EL score”). As expected, Phylonet obtained the lowest EL scores under all model conditions since it estimates species trees by minimizing extra lineages (resulting from deep coalescence). However, the true species trees may have higher amounts of extra lineages as we can see in Fig. 4. ASTRAL, in general, overestimates the EL scores. Overall, these results indicate that Phylonet underestimates the amount of ILS by estimating trees that minimize the number of extra lineages.

**Figure 4:**
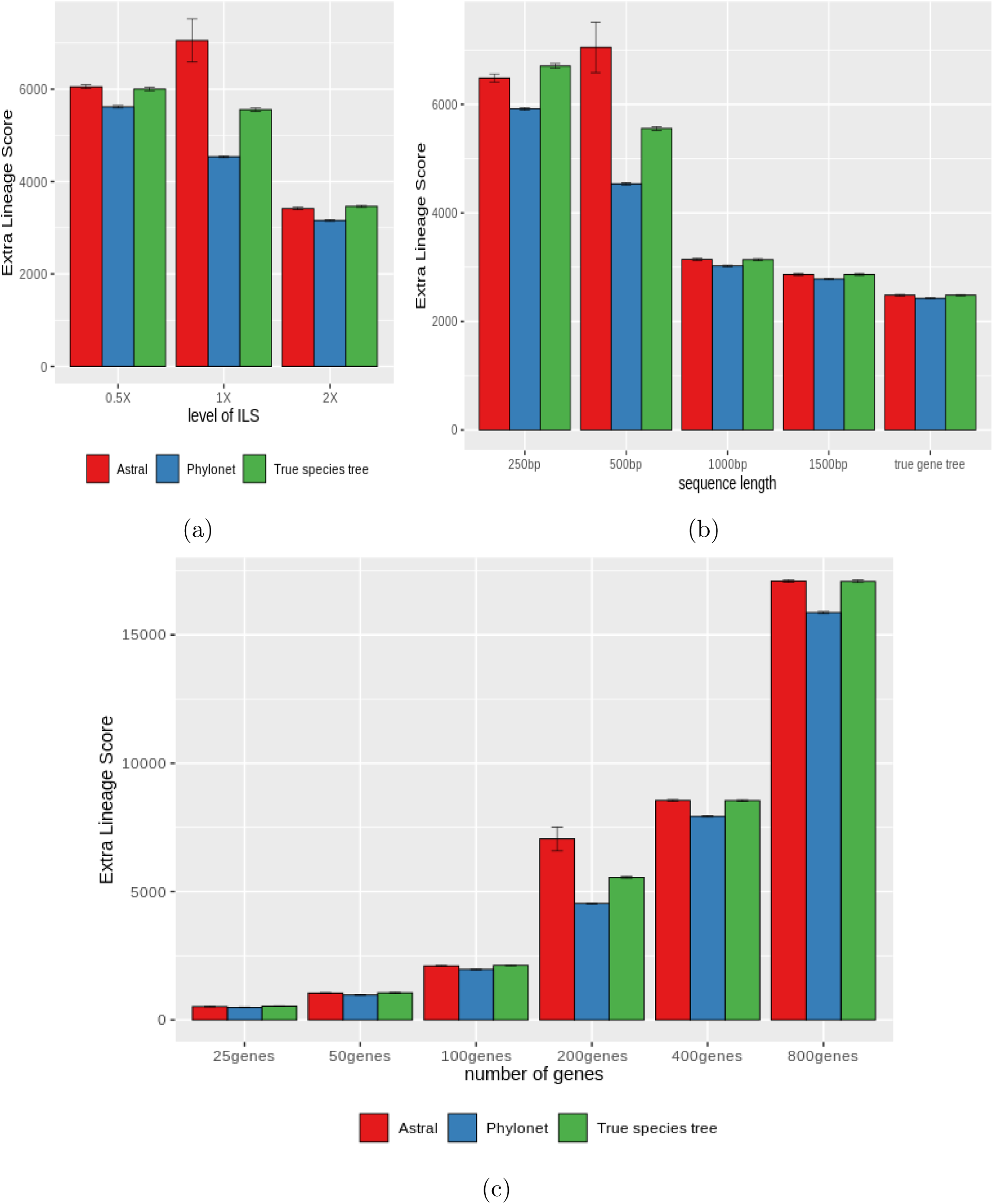
EL scores for ASTRAL, Phylonet and true speciestree on biologically based simulated 37-taxon mammalian dataset. We show average EL scores with standard error bars over 20 replicates for various model conditions (controlling the levels of ILS, gene tree estimation error and numbers of genes).

The EL scores of ASTRAL estimated trees are much closer to the true EL score compared to Phylonet. This is due to the fact that topological accuracy of ASTRAL is much higher than Phylonet and hence embedding/reconciling the gene trees inside the ASTRAL estimated species trees results into closer EL scores (with respect to the true EL scores).

#### Phylogenomic terraces

We have observed that Phylonet may achieve as high quartet score as ASTRAL but may not be as accurate as ASTRAL. That means we may have a collection of trees with the same quartet score but with different topologies – indicating the presence of quartet terraces. To further investigate this, we generated about 9500 neighboring species trees of the species trees estimated by ASTRAL and Phylonet (on a single replicate in 1X, 200g, 500bp model condition) using subtree prune- and-regraft (SPR) operations. We plotted the QS scores of these trees against their corresponding FN rates (see Fig. 5 (a)). This figure clearly shows the presence of various clusters of trees (trees corresponding to the points that lie on a horizontal line) with similar quartet scores but different FN rates, as well as the presence of various clusters of trees (trees corresponding to the points that lie on a vertical line) with the same FN rate but with different quartet scores. Surprisingly, we can observe substantial differences in the topologies of the trees in a quartet terrace (for example, the tree corresponding to the leftmost point on a horizontal line is significantly different than the rightmost tree). Similarly, there are substantial differences in quartet scores of the trees in a cluster of trees with the same FN rate (for example, the trees corresponding to the points at the bottom are substantially different than those at top in a vertical line in terms of the QS). These results support previous findings reported in [43], where they observed some cases where wQMC [30] produced trees with better quartet support scores than ASTRAL but ASTRAL matched the accuracy of wQMC.

**Figure 5:**
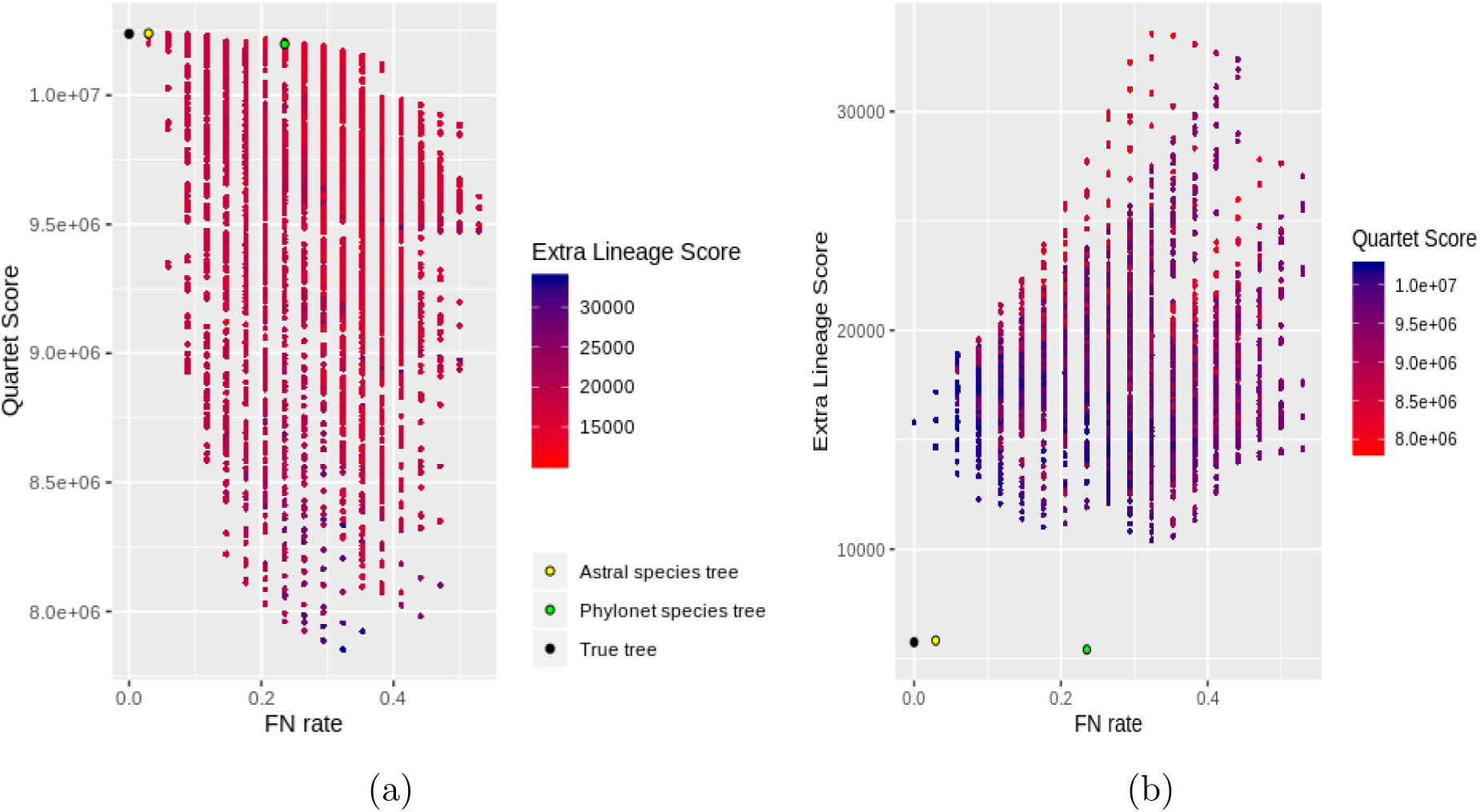
Demonstration of quartet- and MDC-terrace in 37-taxon dataset. We show the results for around 9500 trees: around 4700 neighbors of both ASTRAL- and Phylonet-estimated trees were generated using subtree prune-and-regraft (SPR) operations. We also show the scores of the true species tree. (a) Species tree estimation error vs. quartet score for ASTRAL- and Phylonet-estimated trees and their neighboring trees. (b) species tree estimation error vs. EL score for ASTRAL- and Phylonet-estimated trees and their neighboring trees. We color the data points with a color gradient which varies continuously from dark red to dark blue with increasing EL or quartet scores.

Similar trends hold when we plotted the EL scores against the FN rates (see Fig. 5 (b)). We found clusters of trees with the same EL score but with different topological accuracies and vice versa – indicating the presence of MDC-terrace.

To further investigate the relationships among FN rate, quartet score and EL score, we color the data points (corresponding to the 9500 trees) plotted in Figure 5 with a color gradient which varies continuously from dark red to dark blue with increasing EL scores (in Fig. 5(a)), and with increasing quartet scores (in Fig. 5 (b)). It suggests that higher quartet scores result into lower EL scores, and higher EL scores correspond to lower quartet score. Moreover, the trees with relatively higher FN rates fall somewhat in the middle of the spectrum, meaning that they have moderate quartet and EL scores. From the scores of the ASTRAL- and Phylonet-estimated trees and the true species tree (see the yellow, green and black dots, respectively), it is evident that both ASTRAL and Phylonet “overshoot” respective optimization criteria – ASTRAL *overestimates* the quartet score, and Phylonet *underestimates* the EL score. In doing so, ASTRAL also tends to overestimate the EL score and Phylonet tends to underestimate the quartet score. This underscores the need for applying multi-objective optimization [63, 64] in species tree estimation, where multiple optimization criteria (e.g, MQC and MDC) would be simultaneously optimized to reduce the tendency of overshooting a particular criterion. Similar results are shown separately for the neighborhoods of the ASTRAL- and Phylonet-estimated trees in Figures S7 and S8 in Supplementary Material SM1.

### Results on 11-taxon dataset

11-taxon dataset was simulated under a complex process in order to have substantial heterogeneity between genes and to deviate from the molecular clock [58]. It contains two model conditions: one with long branches that produce low levels of ILS (weak ILS), and the other one with short branches that result into high amounts of ILS (strong ILS). We analyzed the estimated maximum likelihood gene trees as well as the true gene trees. We also varied the numbers of genes from 5 to 100.

#### Missing branch rate

Figure 6 shows the average FN rates of ASTRAL and Phylonet on various model conditions. Both these methods improved as we decreased the amounts of ILS and increased the numbers of genes. Unlike the 37-taxon dataset, under most of the model conditions (strong ILS and weak ILS true gene trees, and weak ILS estimated gene trees), Phylonet matched the accuracy of ASTRAL. ASTRAL improved on Phylonet only on the strong ILS data with estimated gene trees. Both ASTRAL and Phylonet produced highly accurate species trees on true gene trees, especially for weak ILS where they recovered the true species tree even with only 5 genes. Therefore, MDC seems to be more sensitive to the gene tree estimation error than MQC.

**Figure 6:**
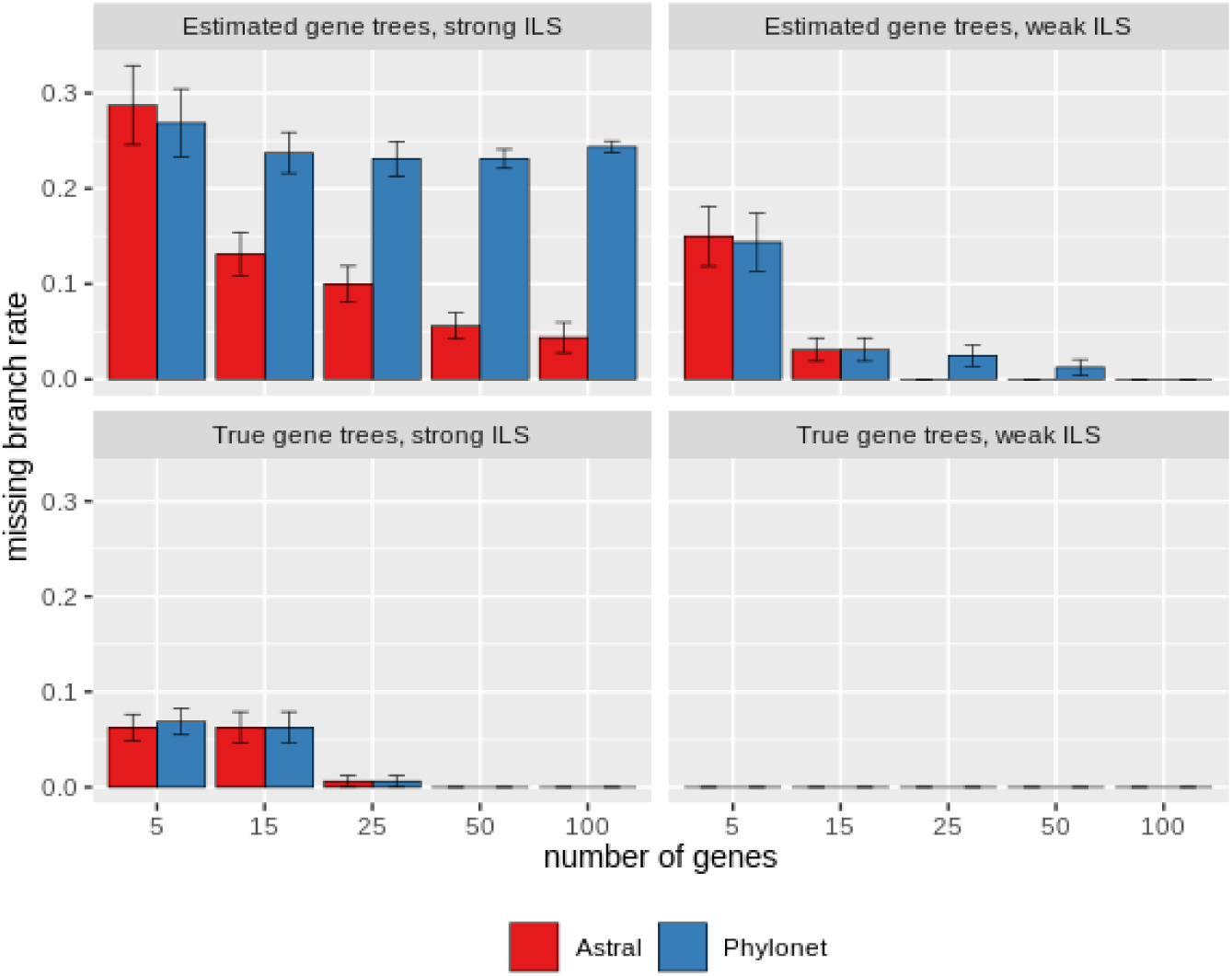
Average FN rates of ASTRAL, and Phylonet on the 11-taxon dataset. We analyzed two model conditions with varying amounts of ILS (strong ILS and weak ILS). We analyzed both estimated and true trees. We varied the numbers of genes from 5 to 100. Average FN rates are shown with standard error bars over 20 replicates.

#### Quartet score

Since both ASTRAL and Phylonet recovered the true species tree on most of the model conditions, the quartet scores are equal to the true QS on those model conditions (see Fig. 7). On the model conditions where they had differences, ASTRAL obtained higher quartet score than Phylonet.

**Figure 7:**
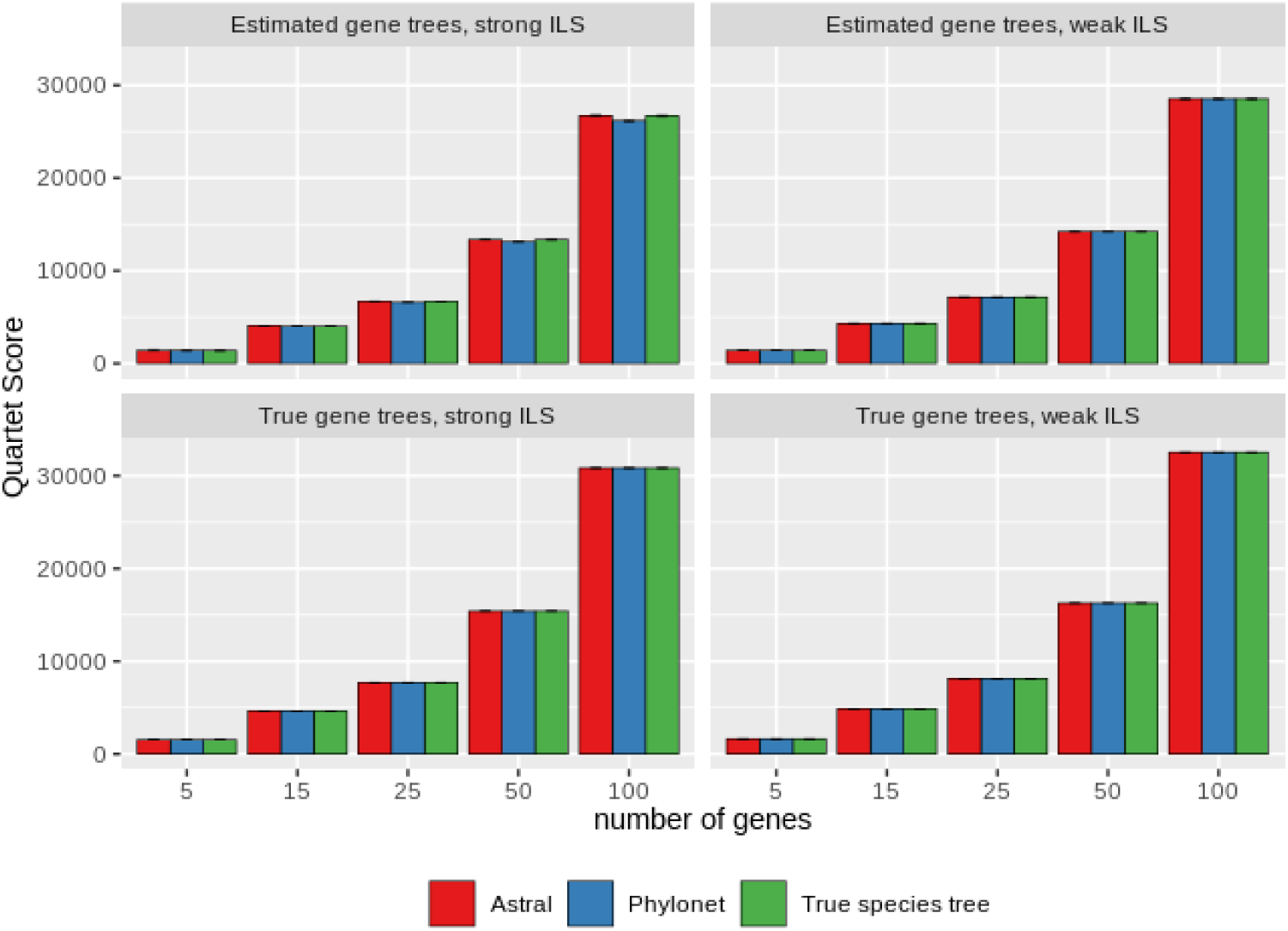
Quartet consistency scores for ASTRAL and Phylonet on 11-taxon dataset under various model conditions. We show average numbers of consistent quartets with standard error bars over 20 replicates.

#### Extra lineage score

Figure 8 shows the EL scores of various trees on 11-taxon dataset. On estimated gene trees, Phylonet achieved significantly lower EL scores than the true EL scores. However, ASTRAL’s EL scores are more closer to the true EL scores. On true gene trees, both Phylonet and ASTRAL recovered the true species tree in most of the cases. But the EL scores of ASTRAL trees are much higher than the true EL scores. This is because ASTRAL returns unrooted trees, and various rooted versions of an unrooted tree have the same FN rate but they do not necessarily have the same EL score (since EL score is sensitive to the rooting of a tree). The search under MDC (by Phylonet) produced the correct rooted versions and hence their EL scores are more closer to true scores than ASTRAL. Computing the EL score of ASTRAL-estimated trees after rooting them on the branch leading to the outgroup will produce lower EL scores as, in most of the cases, ASTRAL-estimated trees on true gene trees are topologically identical to the unrooted version of the model tree.

**Figure 8:**
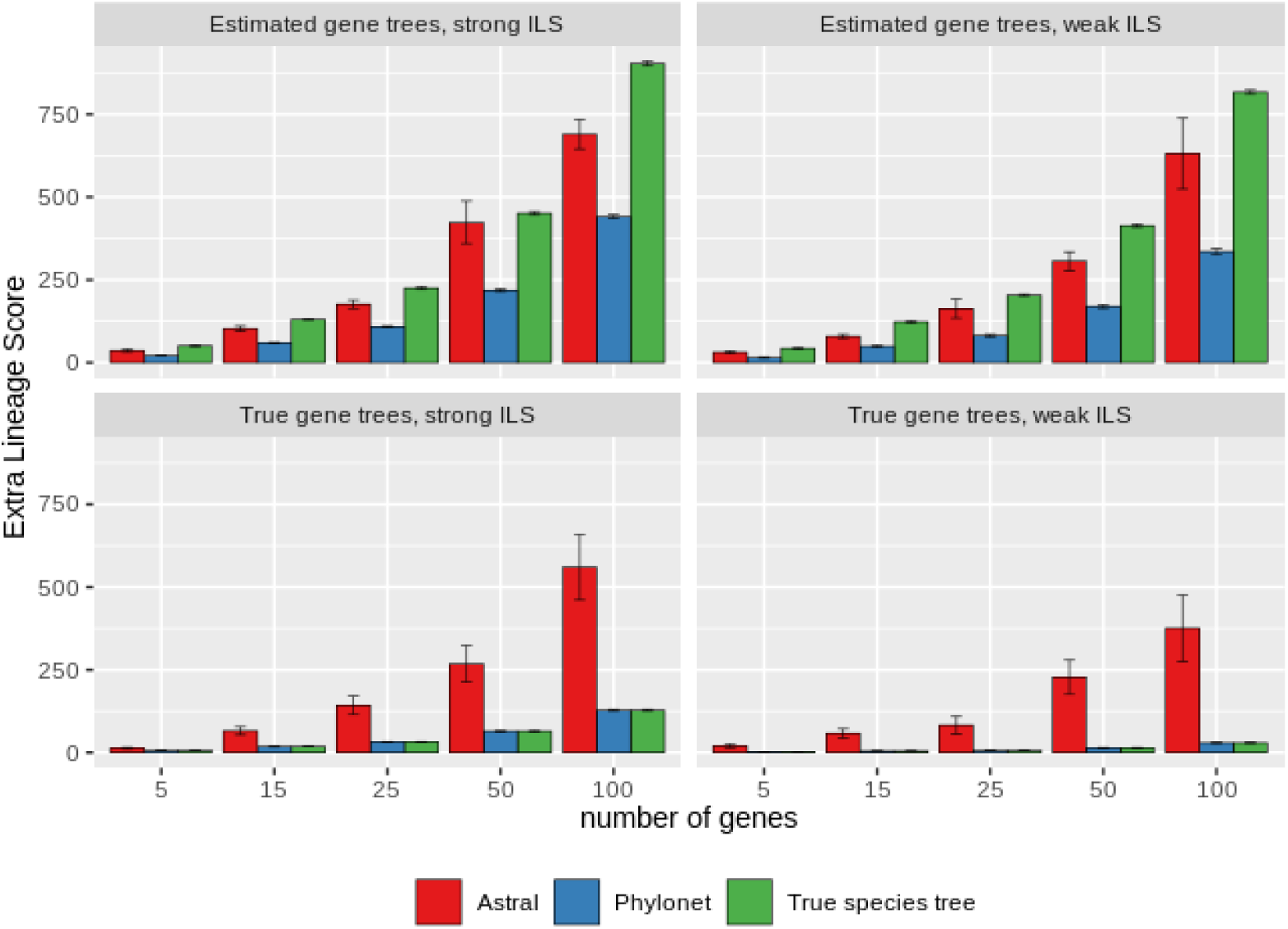
Extra lineage scores for ASTRAL-III and Phylonet on 11-taxon dataset under various model conditions. We show average EL scores with standard error bars over 20 replicates.

#### Phylogenomc terraces

Similar to the 37-taxon dataset, we observed the presence of phylogenomic terraces in 11-taxon dataset. We investigated the FN rates, quartet scores, and EL scores of the trees in the neighborhood of ASTRAL- and Phylonet-estimated trees on a single replicate in strong ILS, 25 gene model condition, and observed MDC and quartet terraces (Figure 9). We found a few replicates (among the 20 replicates) under different model conditions, where Phylonet and ASTRAL obtained identical quartet scores but differ in tree topologies – clearly indicating the presence of quartet-terraces (see Table 3). Similarly, we found some replicates where ASTRAL- and Phylonet-estimated trees are topologically different, having identical EL scores – indicating the presence of MDC-terrace (see Table 4). Similar trends, as in 37-taxon dataset, were observed from the color gradient. Figs. S9 and S10 in Supplementary Material SM1 present the same analyses, but performed separately on the neighborhoods of ASTRAL and Phylonet tree.

**Table 3:** Quartet-terrace on 11-taxon dataset. We show various replicates (among the 20 replicates of data that we analyzed for each of the model conditions), where Phylonet and ASTRAL achieved identical quartet score but differ in tree topologies.

| Model Condition | Replicate No. | Quartet Score |  | FN rate |  |
| --- | --- | --- | --- | --- | --- |
|  |  | Phylonet | Astral | Phylonet | Astral |
| True gene trees,<br>Strong ILS, 5 genes | 2 | 1528 | 1528 | 0.125 | 0 |
|  | 3 | 1536 | 1536 | 0.125 | 0 |
|  | 5 | 1548 | 1548 | 0 | 0.125 |
|  | 10 | 1502 | 1502 | 0 | 0.125 |
|  | 12 | 1548 | 1548 | 0.125 | 0 |
|  | 16 | 1518 | 1518 | 0 | 0.125 |
|  | 20 | 1488 | 1488 | 0.125 | 0 |
| Estimated gene trees,<br>Strong ILS, 5 genes | 19 | 1408 | 1408 | 0.125 | 0 |
| Estimated gene trees,<br>Weak ILS, 5 genes | 12 | 1502 | 1502 | 0.125 | 0.25 |
|  | 15 | 1534 | 1534 | 0.125 | 0.25 |

**Table 4:** MDC-terrace on 11-taxon dataset. We show various replicates (among the 20 replicates of data we analyzed for each of the model conditions), where Phylonet and ASTRAL achieved identical EL scores but differ in tree topologies.

| Model Condition | Replicate No. | EL Score |  | FN rate |  |
| --- | --- | --- | --- | --- | --- |
|  |  | Phylonet | Astral | Phylonet | Astral |
| True gene trees,<br>Strong ILS, 5 genes | 5 | 7 | 7 | 0 | 0.125 |
|  | 10 | 10 | 10 | 0 | 0.125 |
|  | 12 | 6 | 6 | 0.125 | 0 |
|  | 16 | 7 | 7 | 0 | 0.125 |
|  | 20 | 8 | 8 | 0.125 | 0 |

**Figure 9:**
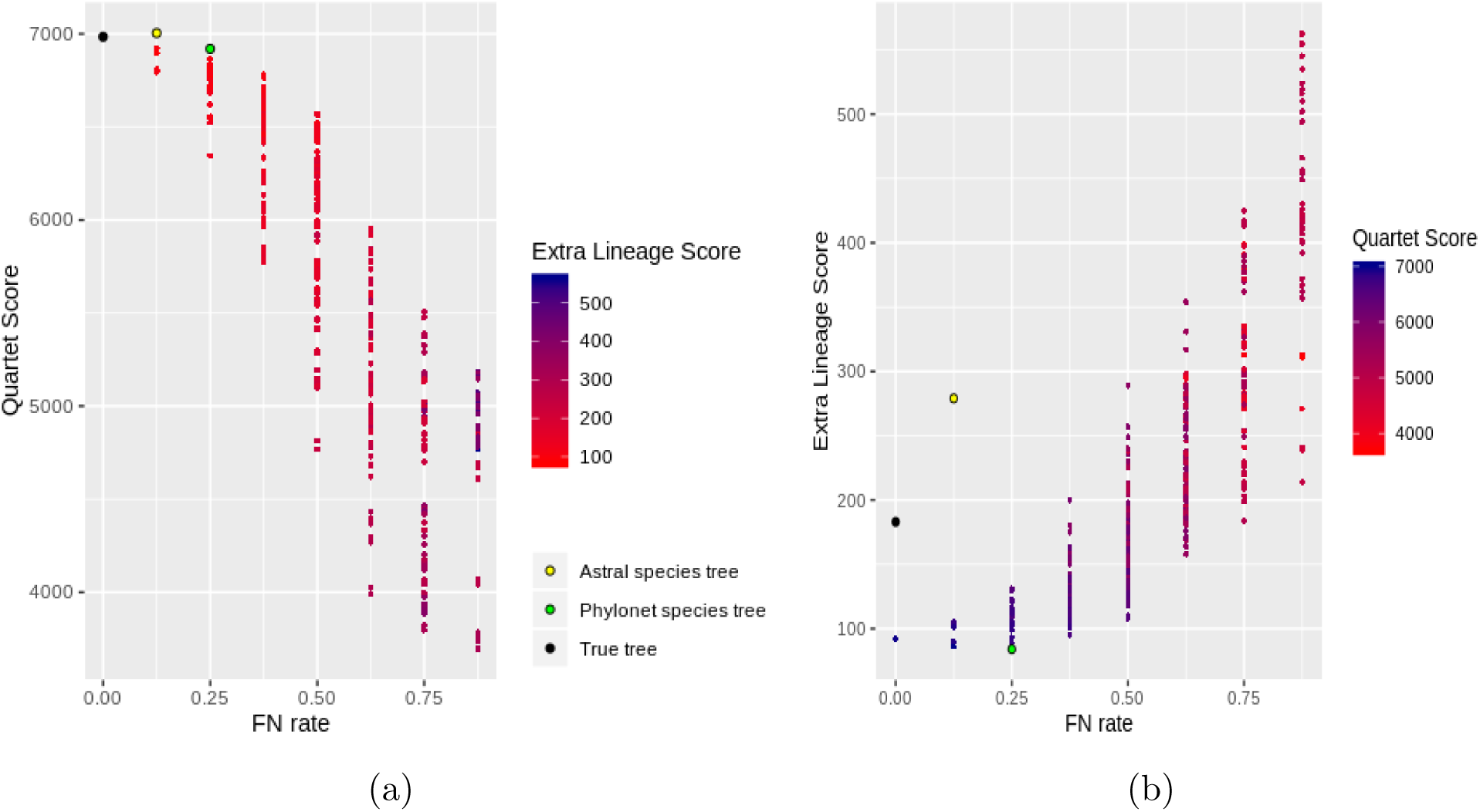
Demonstration of quartet- and MDC-terrace in 11-taxon dataset. We show the results for around 500 trees: around 250 neighbors of both ASTRAL- and Phylonet-estimated trees were generated using subtree prune-and-regraft (SPR) operations. We also show the scores of the true species tree. (a) Species tree estimation error vs. quartet score for ASTRAL- and Phylonet-estimated trees and their neighboring trees. (b) species tree estimation error vs. EL score for ASTRAL- and Phylonet-estimated trees and their neighboring trees. We color the data points with a color gradient which varies continuously from dark red to dark blue with increasing EL or quartet scores.

### Results on 10- and 15-taxon dataset

We used a 15-taxon dataset (previously analyzed in [60]) which was simulated under a model species tree with a caterpillar-like topology, which has 12 short internal branches (0.1 in coalescence units) in succession, a condition that gives rise to high levels of ILS [1, 11]. There are four model conditions with varying gene sequence lengths (100 or 1000 sites), and varying numbers of genes (100 and 1000). We also analyzed a 10-taxon dataset where there is a different species tree for each replicate, and 200 gene trees were simulated for each species tree using the multi-species coalescent process [60]. 10-taxon dataset has two model conditions: one with very high ILS and another with relatively lower ILS.

Similar trends, as we have observed on 37- and 11-taxond dataset, were observed on 10- and 15-taxon dataset. We present the results on these two dataset in Supplementary Materials SM1.

### Results on biological dataset

#### Amniota dataset

We analyzed the Amniota dataset from Chiari *et al.* [61] containing 248 genes across 16 amniota taxa in order to resolve the position of turtles relative to birds and crocodiles. This dataset contains both amino acid (AA) and nucleotide (DNA) gene trees. We ran exact versions for both ASTRAL and Phylonet-MDC.

Previous studies suggest that placing turtles as a sister group to Archosaurs (birds and turtles) is more reliable [61, 65]. Astral on both DNA and AA data produced (birds,(turtles,crocodiles)), and thus recovered the Archosaurs (see Fig. 10). Phylonet estimated trees on both AA and DNA gene trees are same except for the resolution within the turtles (*Phrynops hilarii, Caretta caretta, Chelonoidis nigra, and Emys orbicularis*). Phylonet, on both AA and DNA data, produced the Archosaurs, but it placed turtles as sister to Squamates (snakes and lizards) and placed Archosaurs as sister to the clade containing turtles and Squamates. Thus, phylonet did not produce the (birds,(turtles,crocodiles)) relationship. The quartet and extra lineage scores of ASTRAL and Phylonet-MDC estimated trees are given in Tables 5.

**Table 5:** Quartet and EL scores of ASTRAL- and Phylonet-estimated trees on the amniota dataset (both DNA and AA).

| Tool | EL score | Quartet score |
| --- | --- | --- |
| ASTRAL (DNA) | 2620 | 97890 |
| Phylonet (DNA) | 1170 | 93018 |
| ASTRAL (AA) | 3293 | 83604 |
| Phylonet (AA) | 1916 | 80507 |

**Figure 10:**
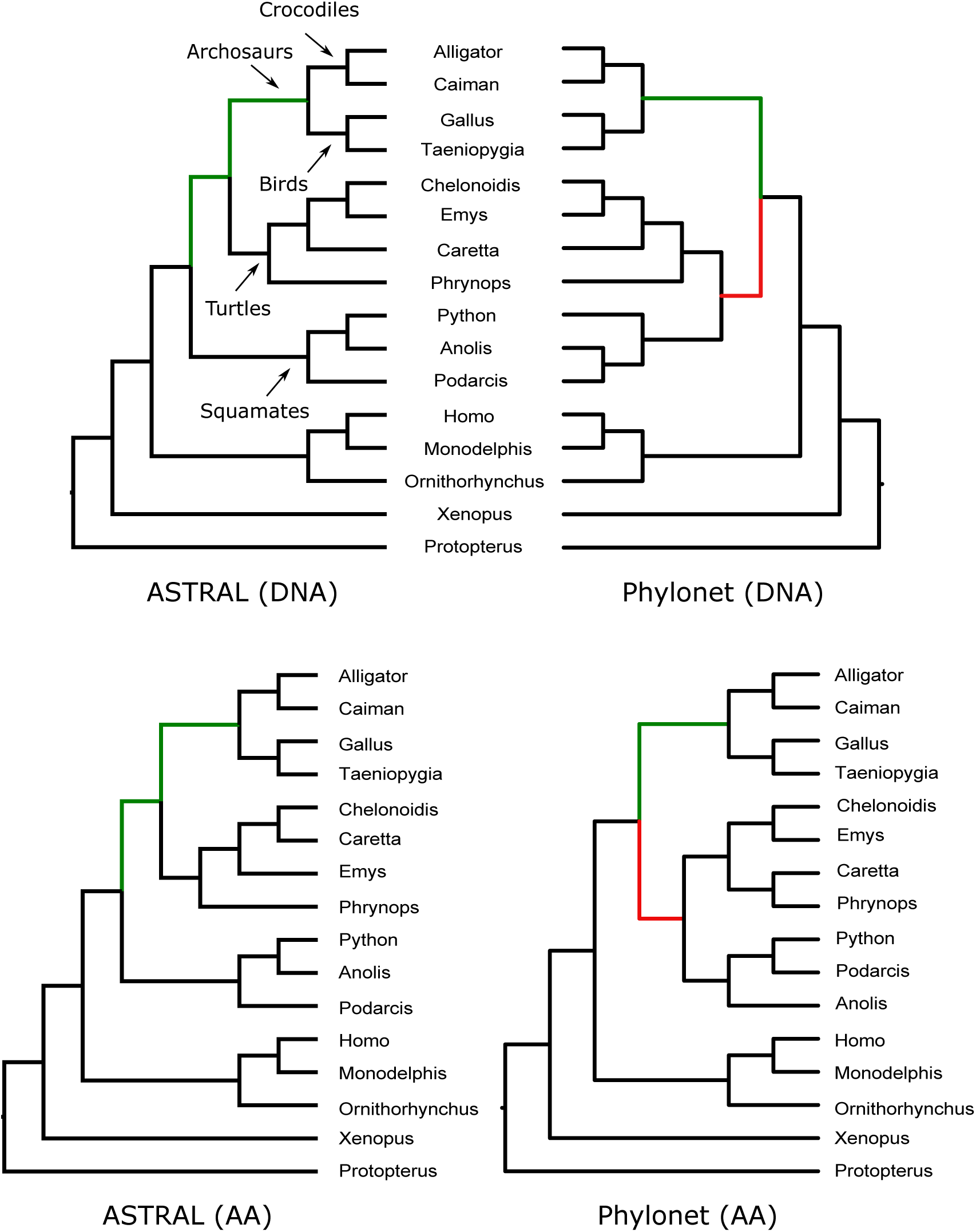
Analyses of the Amniota dataset (both DNA and AA gene trees) by maximizing quartet score (ASTRAL) and minimizing extra lineage score (Phylonet). We show the rooted versions of the ASTRAL-estimated trees using the outgroup (*Protopterus annectens*).

#### Mammalian dataset

Song *et al.* analyzed a dataset containing 447 genes across 37 mammals using MP-EST [12] and combined analyses (CA-ML) [59]. We reanalyzed this dataset after removing 21 mislabeled genes (confirmed by the authors), and two other outlier genes using ASTRAL and Phylonet. The placement of bats (*Myotis lucifugus* and *Pteropus vampyrus*) and tree shrews (*Tupaia belangeri*) were two of the questions of greatest interest, and alternative relationships have previously been reported [66–69]. ASTRAL and phylonet estimated trees also exhibit similar differences with respect to the placement of bats and tree shrews (see Fig. 11). ASTRAL placed tree shrews as sister to Glires (Rodentia, Lagomorpha) which is consistent to the tree estimated by combined analysis using maximum likelihood (CA-ML) reported in [59]. Phylonet recovered a tree that placed tree shrews as sister to the Primates, which is consistent to the tree estimated by MP-EST using multi-locus bootstrapping [70] (reported in [13, 59, 65]). Both trees put Perissodactyla (*Equus caballus*) as a sister to Carnivora (*Canis familiaris, Fellis catus*). With respect to the position of bats, ASTRAL agrees with MP-EST which placed bats as sister to the (Cetartiodactyla, (Perissodactyla, Carnivora)) clade. Phylonet placed bats as sister to Cetartiodactyla, and put (Perissodactyla, Carnivora) as the sister clade of (bats, Cetartiodactyla), and thus agrees with CA-ML [65]. The extra lineage and quartet scores of these two trees are reported in Table 6.

**Table 6:** Quartet and EL scores of ASTRAL- and Phylonet-estimated trees on the biological mammalian dataset [59].

| Tool | EL score | Quartet score |
| --- | --- | --- |
| ASTRAL | 5909 | 25526915 |
| Phylonet | 5675 | 25479405 |

**Figure 11:**
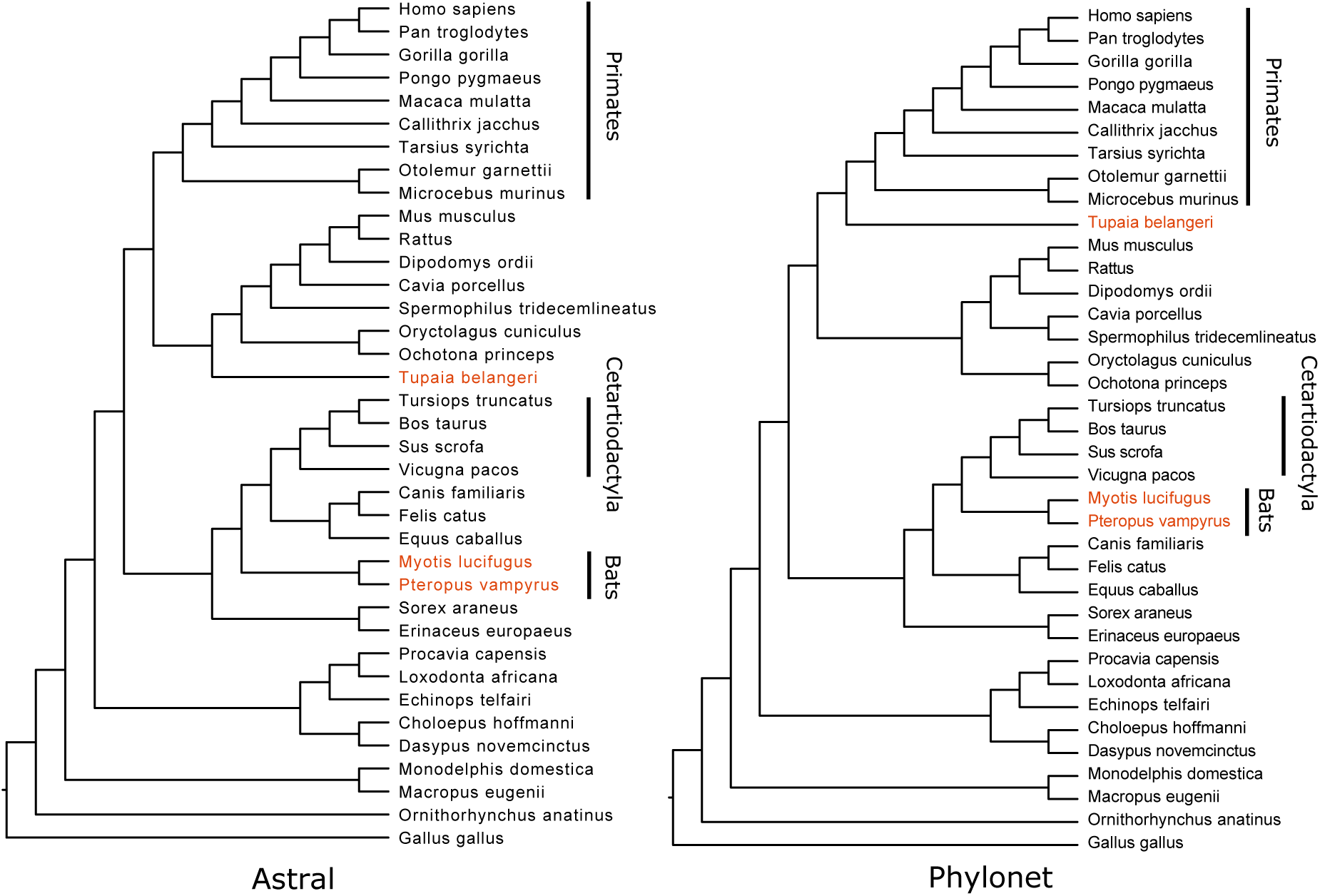
Analyses of the mammalian dataset by maximizing quartet score (ASTRAL) and minimizing deep coalescence (Phylonet). These two trees differ in the placement of tree shrews and bats.

## Conclusions

In this paper, we report, on an extensive evaluation study, the comparison between MQC and MDC criteria for estimating species trees in the presence of incomplete lineage sorting. While the superiority of MQC over MDC is expected (since MQC is a statistically consistent estimator of species tree under ILS), and the observations that MDC criteria underestimates the amount of deep coalescence is not novel [35], this study is the first to evaluate MQC and MDC and confirms these results on various simulated and real biological datasets, and hence provide additional support for the consistency properties of MQC and MDC. Although the presence of multiple equally good species trees with respect to a particular optimization criteria is not unexpected [31], we – for the first time– formalized the concept of phylogenomic terraces when species trees are estimated from a collection of gene trees, and analyzed its impact in the context of MQC and MDC. Our experimental study suggests that species trees estimated under MQC and MDC may fall within terraces and their neighborhoods, but ASTRAL’s search strategy leads to trees within terraces that are closer to the true species trees than the trees estimated under MDC. This study reveals various interesting trends regarding the FN rates, quartet scores and EL scores of the trees estimated by MQC and MDC under various model conditions.

Phylogenomic terraces have implications in the search strategies under various optimization criteria. Species tree space grows exponentially with the number of taxa. Thus, algorithms to find optimal species trees under various optimization criteria requires navigating a large tree space. Considering the presence of large phylogenomic terraces, efficient algorithms to strategically explore the terraces and their neighborhood is crucial. Fundamental to most of the summary methods is the ability to efficiently explore and score (with respect to a particular optimization criterion) the trees inside the tree space. Since all the trees inside a particular terrace have the same optimization score, identifying a terrace may help reduce the computational efforts by avoiding the computation time that might be unnecessarily spent to evaluate many trees with identical score. Thus, efficient identification of a terrace and directing the tree search from one terrace to the other ones with higher optimization scores may result into faster convergence. Exact solutions to various optimization criteria, such as MQC [13], MDC [32] and MGDL [55, 56] are available that are guaranteed to find the globally optimal solution under respective optimization criteria. However, the application of exact versions to large datasets has been limited by the prohibitive amount of time required by the available algorithms to explore the tree space exhaustively. Therefore, utilizing the knowledge of terraces may help prune the search space. However, it could also be possible that a particular tree in a terrace is topologically more correct than the other ones, and hence navigating off from a terrace may lead us to miss more reliable (in terms of topological accuracy) trees. Therefore, the presence of potentially large terraces imposes the challenge of identifying relatively more reliable trees within the terraces and their neighborhoods. Thus, the multiplicity of equally good trees in a species tree terrace introduces ambiguity. One plausible option for reducing the ambiguity is to estimating consensus trees (greedy consensus, majority consensus, maximum agreement subtree, maximum clade credibility tree, etc.) of the trees in a terrace. This may lead us to topologically more accurate trees, and can be used to draw branch supports on the species tree without having to rely on multi-locus bootstrapping [70]. Thus, future studies need to investigate the properties of the consensus trees of the trees in a phylongenomic terrace. Another important direction would be optimizing multiple optimization criteria [63, 64] simultaneously instead of a single one.

Navigating trees within a terrace could be easier due to their similarity with respect to a particular optimization criterion. *terrace-aware* data structures led to substantial speedup of RAxML [71] and IQ-tree [72] for estimating ML trees from alignments [48]. Thus, efficient *terrace-aware* algorithms and data structures for strategically navigating trees both inside a phylogenomic terrace and its neighborhood would contribute to the improvement of the summary methods both in terms of accuracy and scalability. Efficiently characterizing a terrace – by quantifying the difference between the pair of trees using various distance measures such as, average Robinson-Foulds (RF), nearest neighbor interchange (NNI), subtree prune and regraft (SPR), and tree bisection and re-connection (TBR) distances – would be another interesting research direction. Thus, the discovery of terraces poses various challenges as well as opens up several important research avenues.

This study is limited in scope and can be extended in several directions. We analyzed complete gene trees with full set of taxa. Future studies need to investigate the impact of the presence of missing taxa (i.e., incomplete gene trees) in phylogenomic terraces, as missing taxa may hamper the tree search due to the presence of terraces as suggested by previous studies [46, 73]. This study analyzed small to moderate sized dataset. Small dataset enabled us to run the exact versions of ASTRAL and Phylonet. However, the impact of terraces in larger dataset with hundreds of taxa need to be investigated as the possibility of the presence of potentially large terraces is high for larger numbers of taxa. This study investigated relatively long sequences (250 ∼ 2000 bp); subsequent studies should investigate the relative performance of MQC and MDC on very short sequences, since recombination-free loci can be very short [74]. Finally, investigating various structural properties of terraces under various optimization criteria is crucial for developing terrace-aware data structures and tree search algorithms.

## Supporting information

**Supplementary Materials SM1:** Additional results and data.

## Supplementary Materials

These supplementary materials present additional results, supplementary figures and tables, and the commands that we used to run various methods.

### 1 Additional Results

#### Results on 15-taxon datasets

ASTRAL is significantly better (*P <<* 0.05) than Phylonet across all the model conditions. The improvement of ASTRAL over Phylonet increases as we increase the number of genes and gene tree accuracy (by increasing the gene sequence lengths). On true gene trees, ASTRAL returned the true species trees whereas Phylonet incurred around 50% average missing branch rate. In terms of the quartet score, ASTRAL consistently achieved higher quartet consistency than Phylonet and is more close to the quartet scores of the true species trees. Phylonet achieved significantly lower extra lineage scores than the ASTRAL-estimated trees and the true species trees. Thus, similar to the observations on 11- and 37-taxon dataset, these results support that MDC criterion underestimates the amount of ILS present in the dataset, as it tries to minimize the amount of deep coalescence. Interestingly, Phylonet-estimated trees have much closer EL scores than the ASTRAL-estimated trees, even though Phylonet is significantly worse than ASTRAL on this model condition. This suggests that overestimating quartet score is more desirable than underestimating the amount of ILS by minimizing EL score, and hence it provides additional support for the statistical inconsistency of MDC criterion [1].

**Figure S1:**
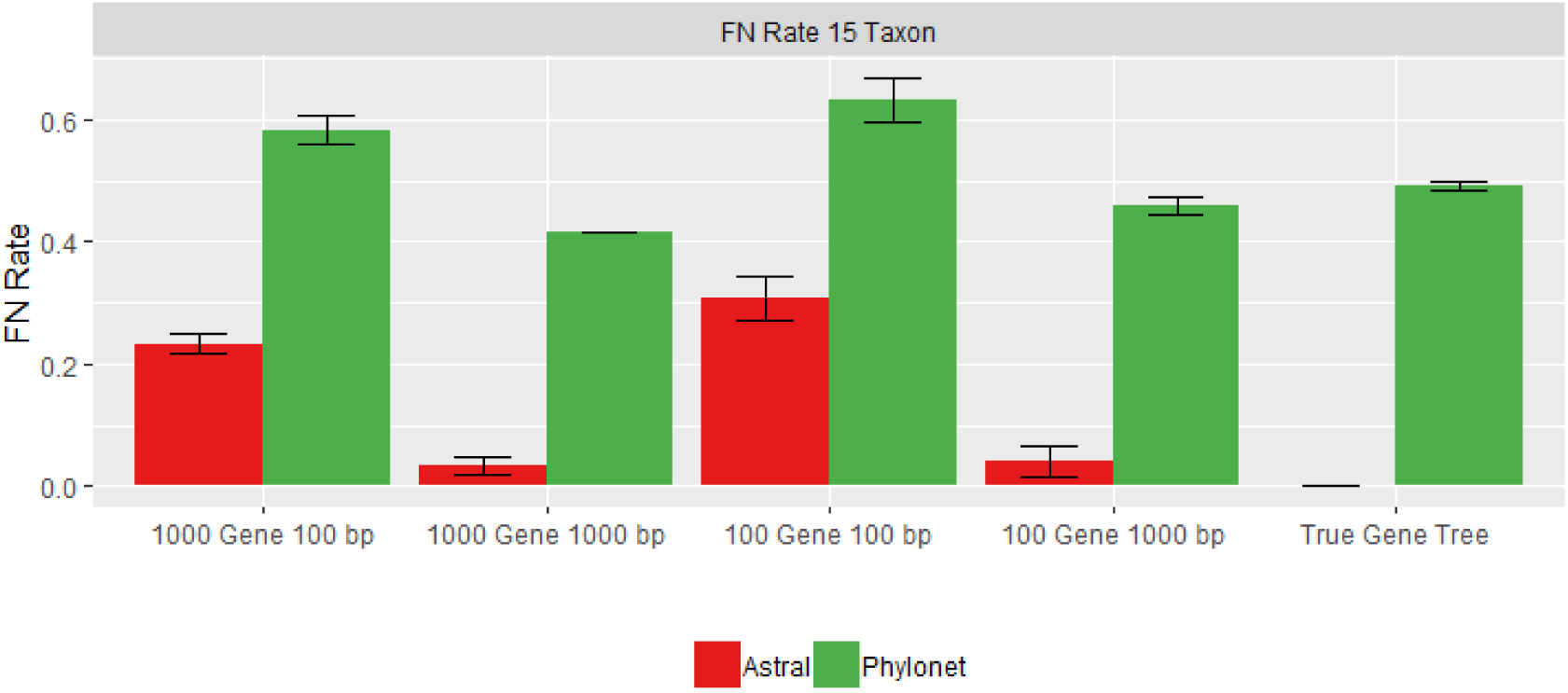
Average missing branch rates of ASTRAL and Phylonet on 15-taxon dataset. Four model conditions on estimated gene trees were analyzed with varying numbers of genes (100 and 1000) and sequence lengths (100 bp and 1000 bp). We also analyzed a model condtions with 1000 true gene trees. We show the average FN rates with standard error bars over 10 replicates.

Another significant trend is that the difference of quartet scores of Phylonet and ASTRAL are not as substantial as the difference in FN rates. Especially, on model conditions with relatively lower and no gene tree estimation error, ASTRAL is significantly better than Phylonet but the differences in quartet scores are not that substantial.

**Figure S2:**
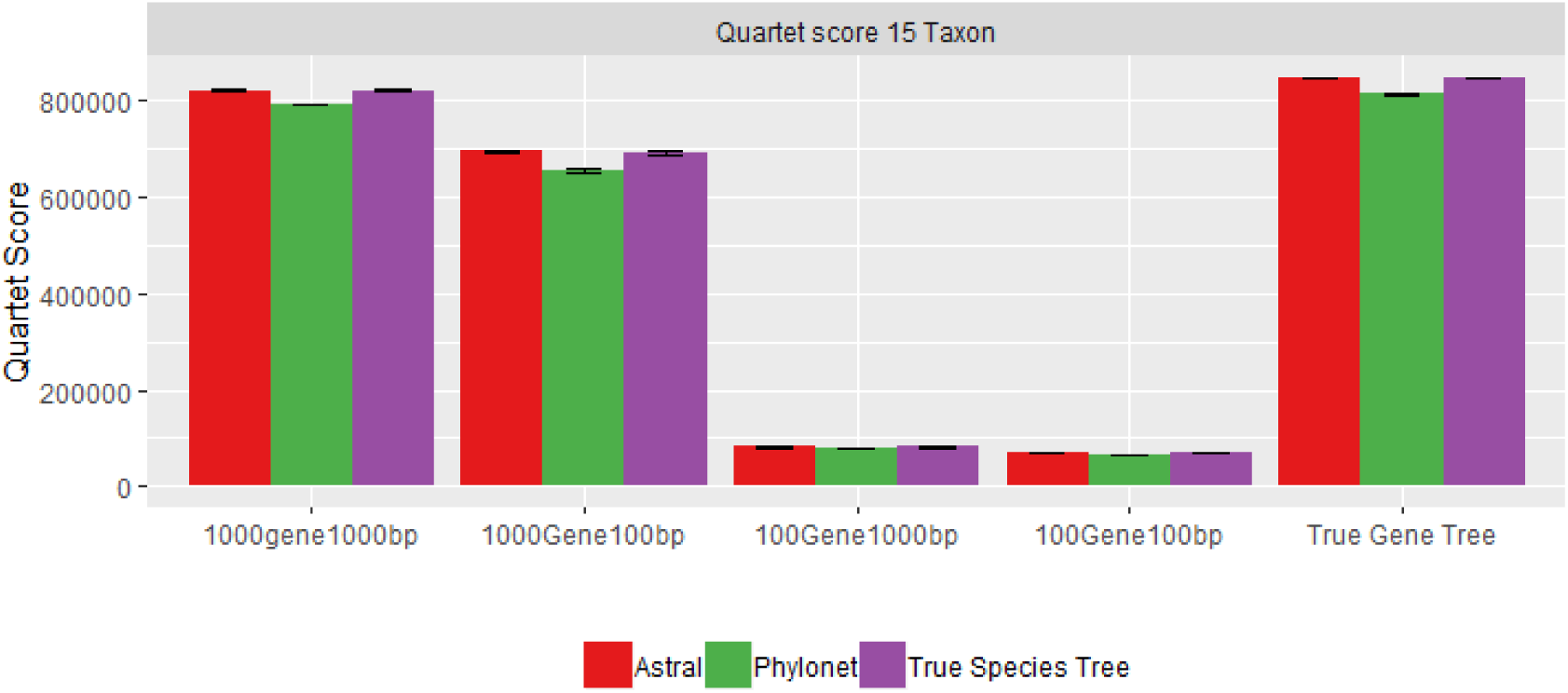
Quartet scores for ASTRAL, Phylonet and true species tree on 15-taxon dataset. We show average quartet scores with standard error bars over 10 replicates for various model conditions.

**Figure S3:**
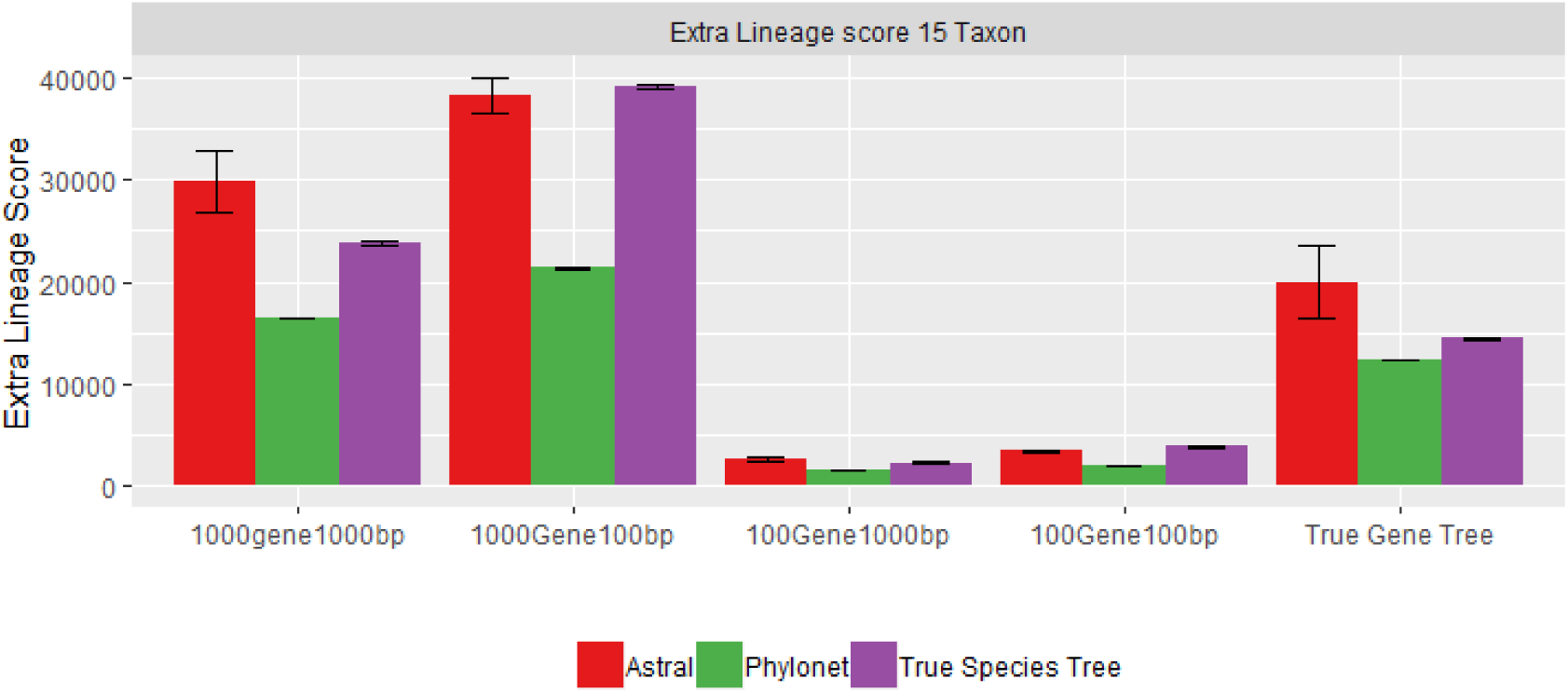
Extra lineage (EL) scores for ASTRAL, Phylonet and true species tree on 15-taxon dataset. We show average EL scores with standard error bars over 10 replicates for various model conditions.

#### Results on 10-taxon datasets

Results on 10-taxon dataset are shown in Figs. S4, S5, and S6.

**Figure S4:**
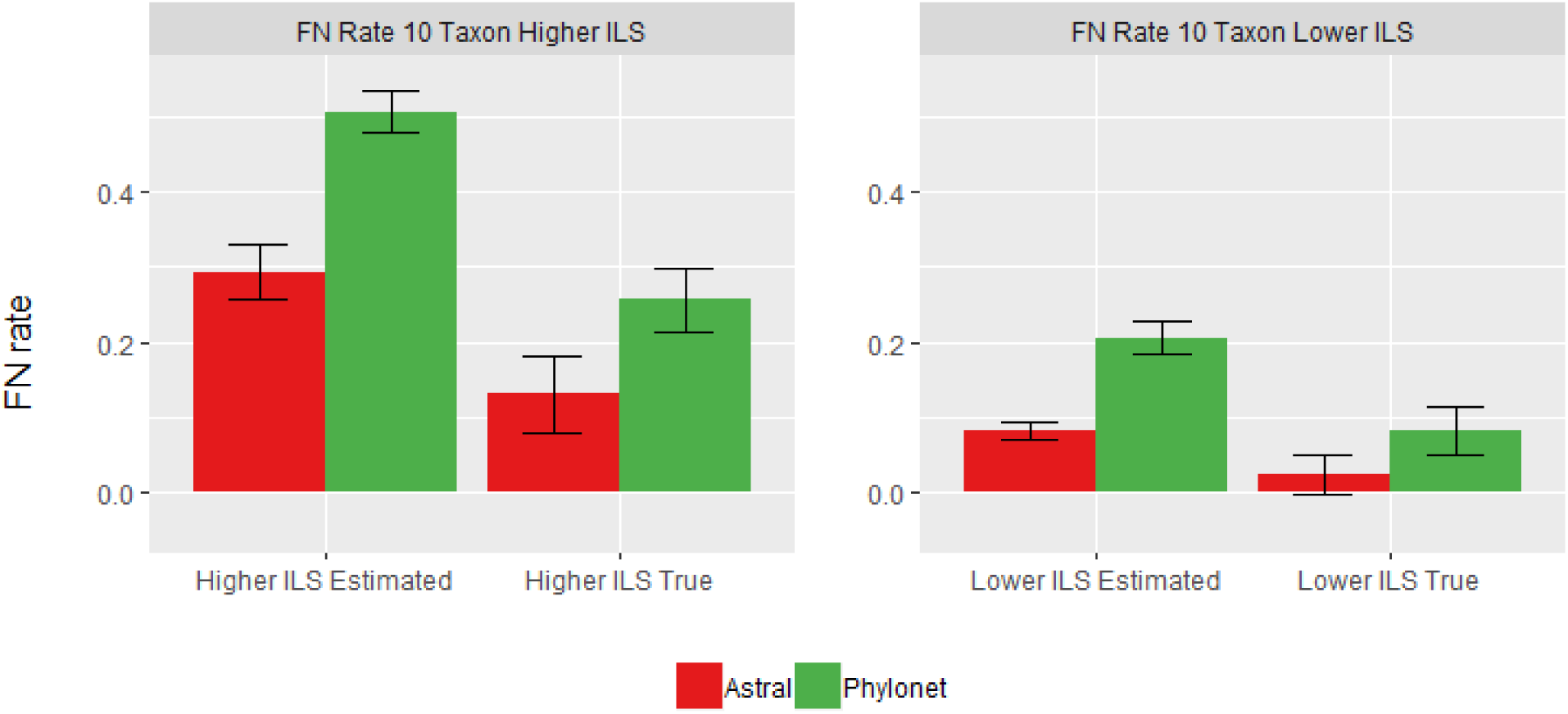
Average missing branch rates of ASTRAL and Phylonet on 10-taxon dataset. We analyzed both estimated and true gene trees with varying amounts of ILS (Higher ILS, and Lower ILS). We show the average FN rates with standard error bars over 20 replicates.

**Figure S5:**
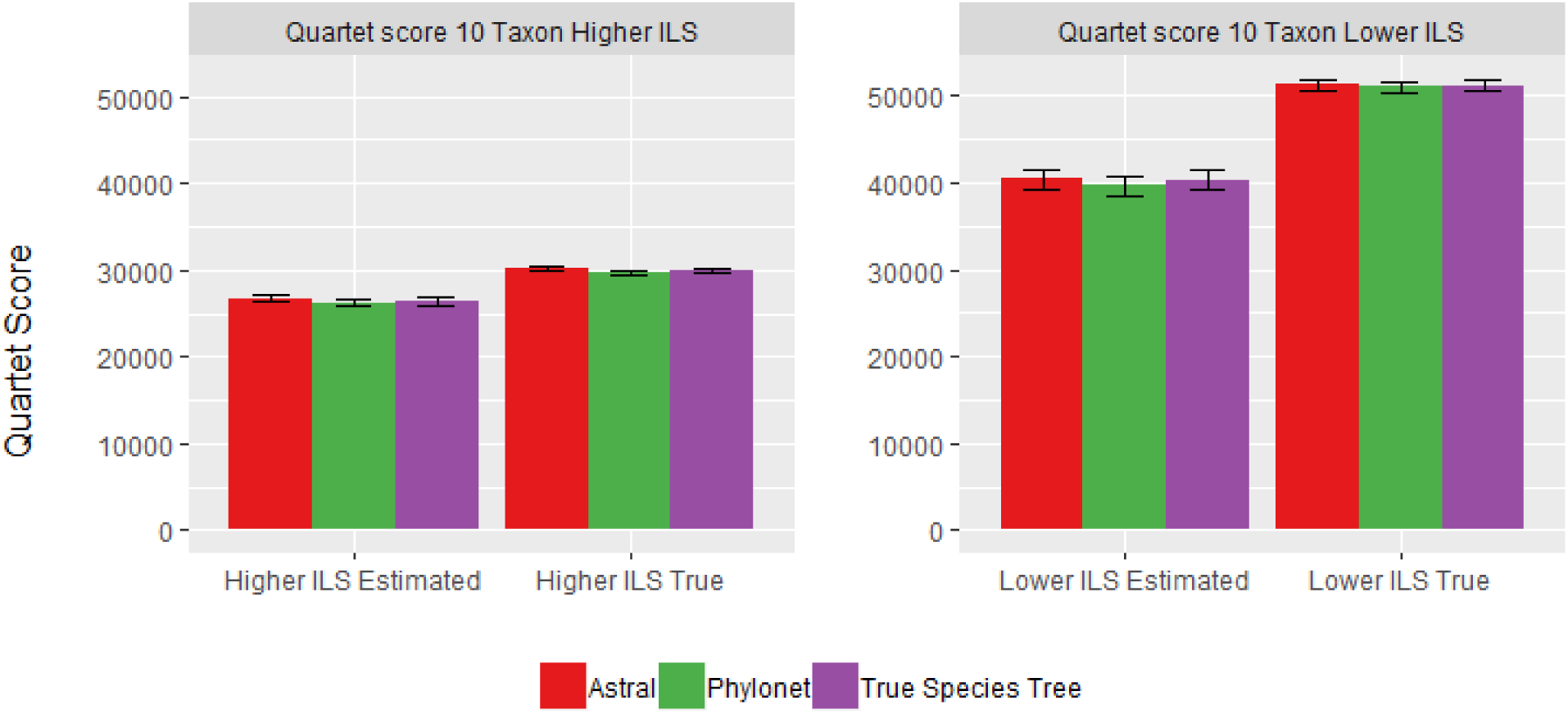
Quartet scores for ASTRAL, Phylonet and true species tree on 10-taxon dataset. We show average quartet scores with standard error bars over 20 replicates for various model conditions.

#### Additional Results on 37-taxon and 11-taxon dataset

**Figure S6:**
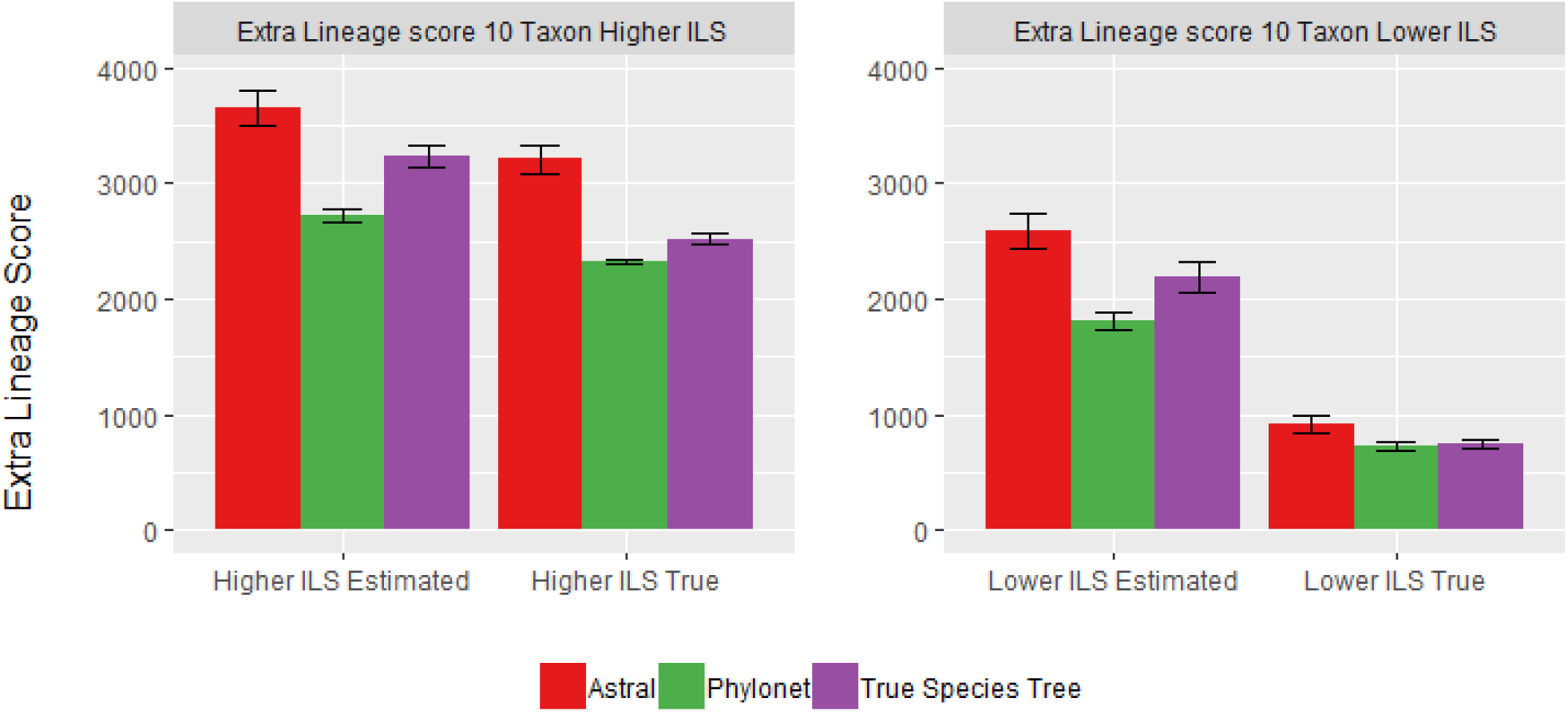
EL scores for ASTRAL, Phylonet and true species tree on 10-taxon dataset. We show average EL scores with standard error bars over 20 replicates for various model conditions.

**Figure S7:**
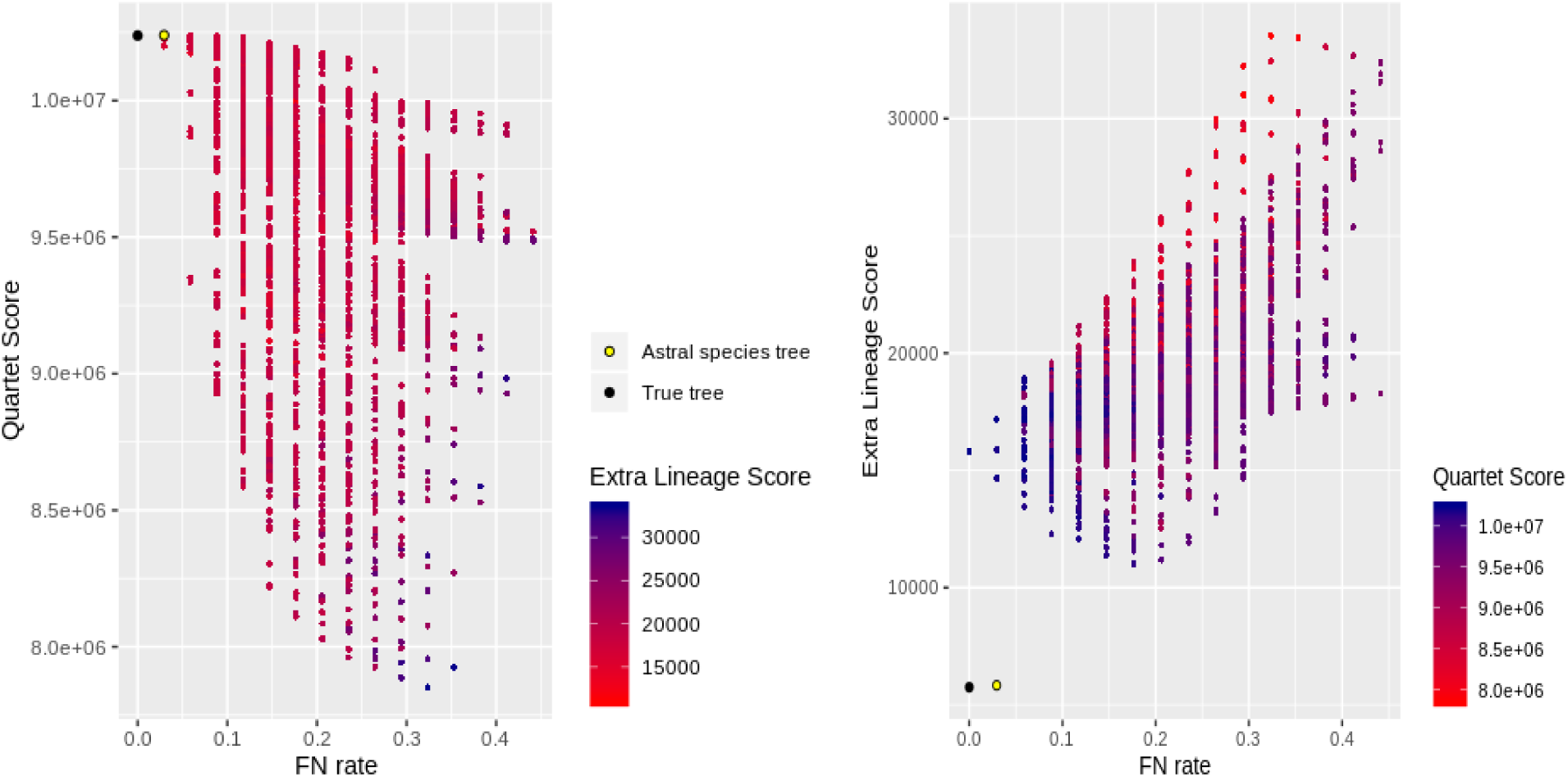
Demonstration of quartet- and MDC-terrace in 37-taxon dataset. We show the results for around 4700 neighboring trees, generated using subtree prune-and-regraft (SPR) operations, of ASTRAL-estimated trees. We also show the scores of the true species tree. (a) Species tree estimation error vs. quartet score for ASTRAL-estimated tree and its neighboring trees. (b) species tree estimation error vs. EL score for ASTRAL–estimated trees and its neighboring trees. We color the data points with a color gradient which varies continuously from dark red to dark blue with increasing EL or quartet scores.

**Figure S8:**
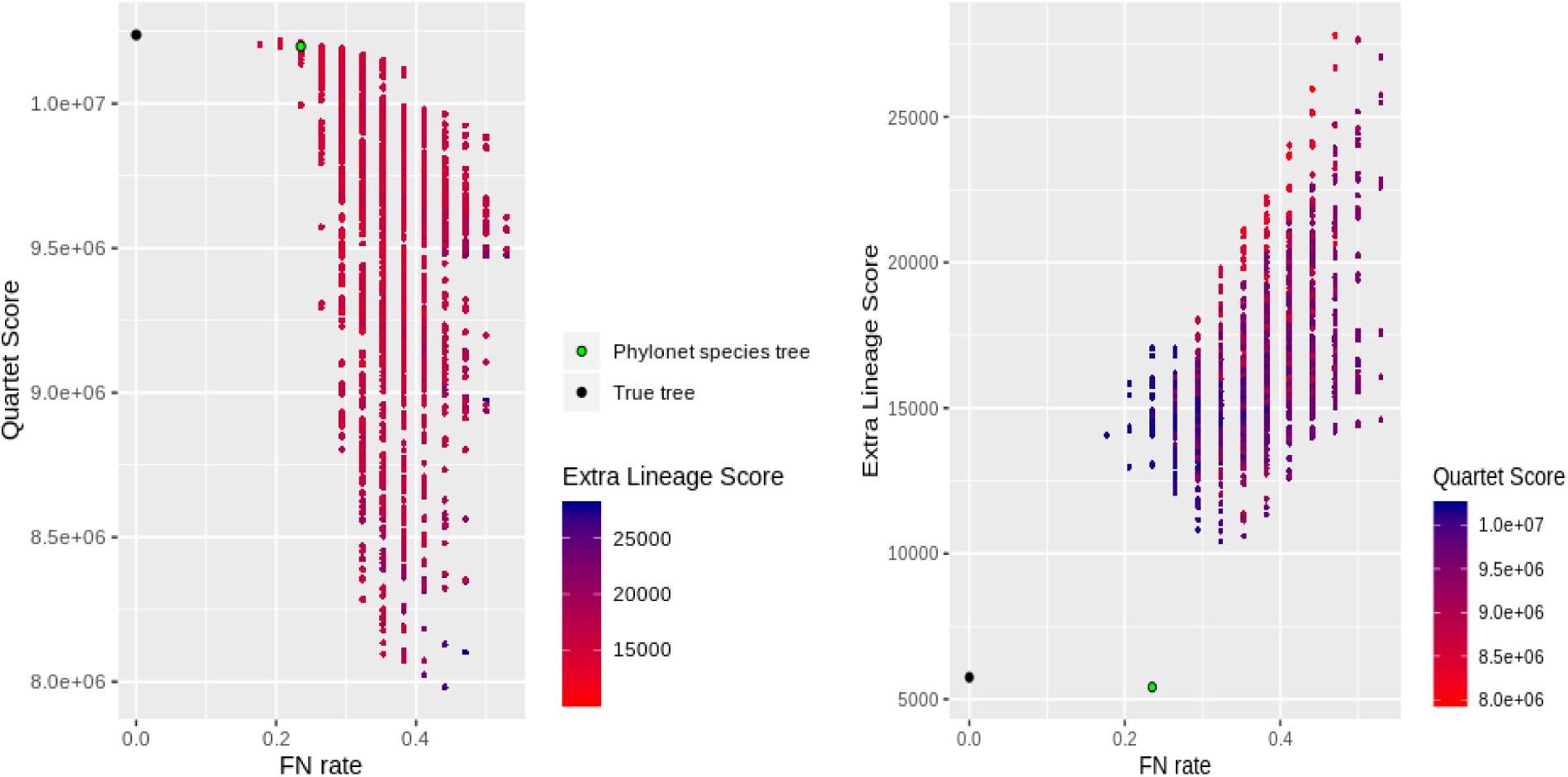
Demonstration of quartet- and MDC-terrace in 37-taxon dataset. Similar to Fig. S7, We show the results for around 4700 neighboring trees of the Phylonet-estimated tree.

**Figure S9:**
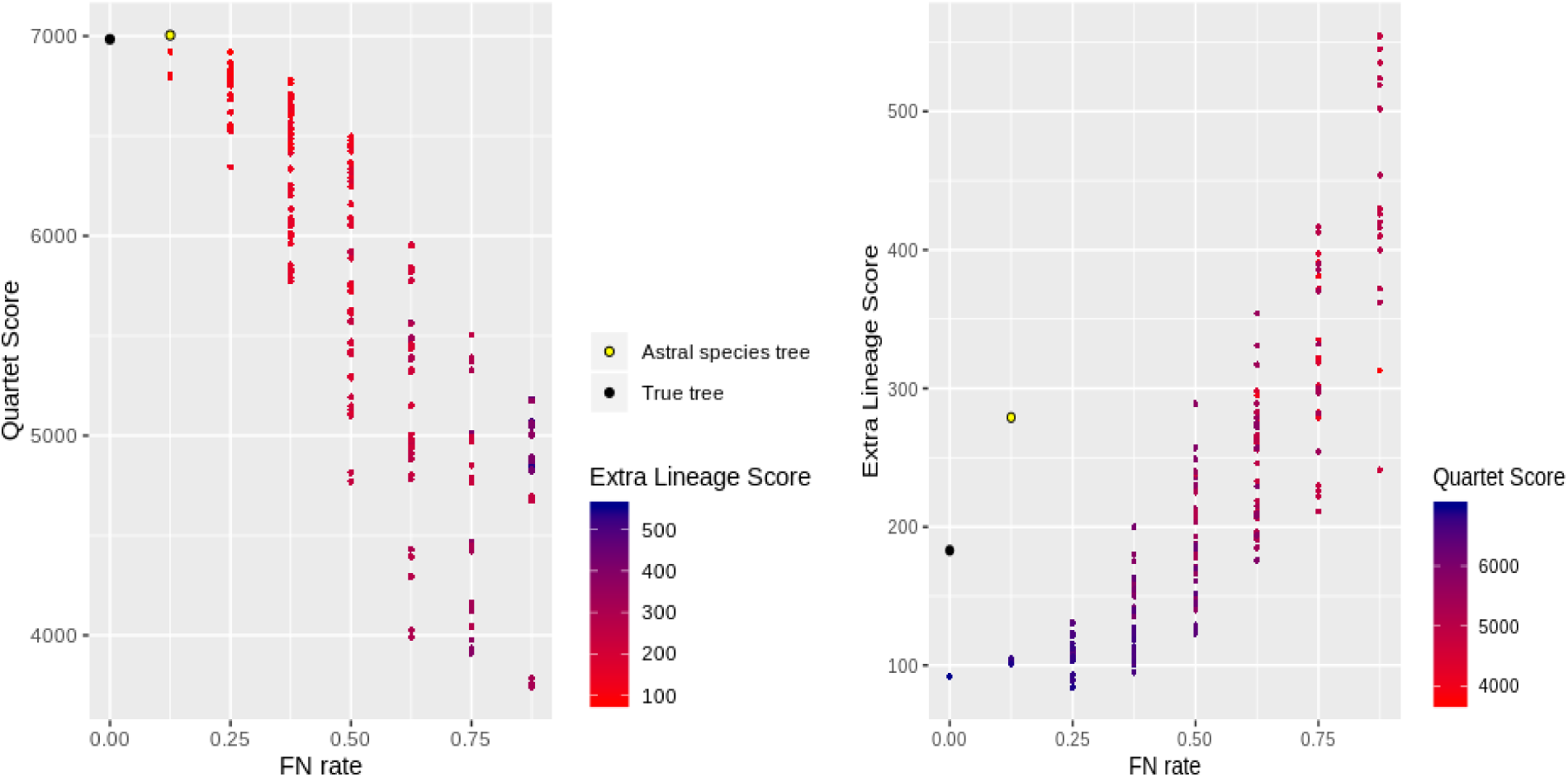
Demonstration of quartet- and MDC-terrace in 11-taxon dataset. We show the results for around 250 neighboring trees, generated using subtree prune-and-regraft (SPR) operations, of ASTRAL-estimated trees. We also show the scores of the true species tree. (a) Species tree estimation error vs. quartet score for ASTRAL-estimated tree and its neighboring trees. (b) species tree estimation error vs. EL score for ASTRAL–estimated trees and its neighboring trees. We color the data points with a color gradient which varies continuously from dark red to dark blue with increasing EL or quartet scores.

**Figure S10:**
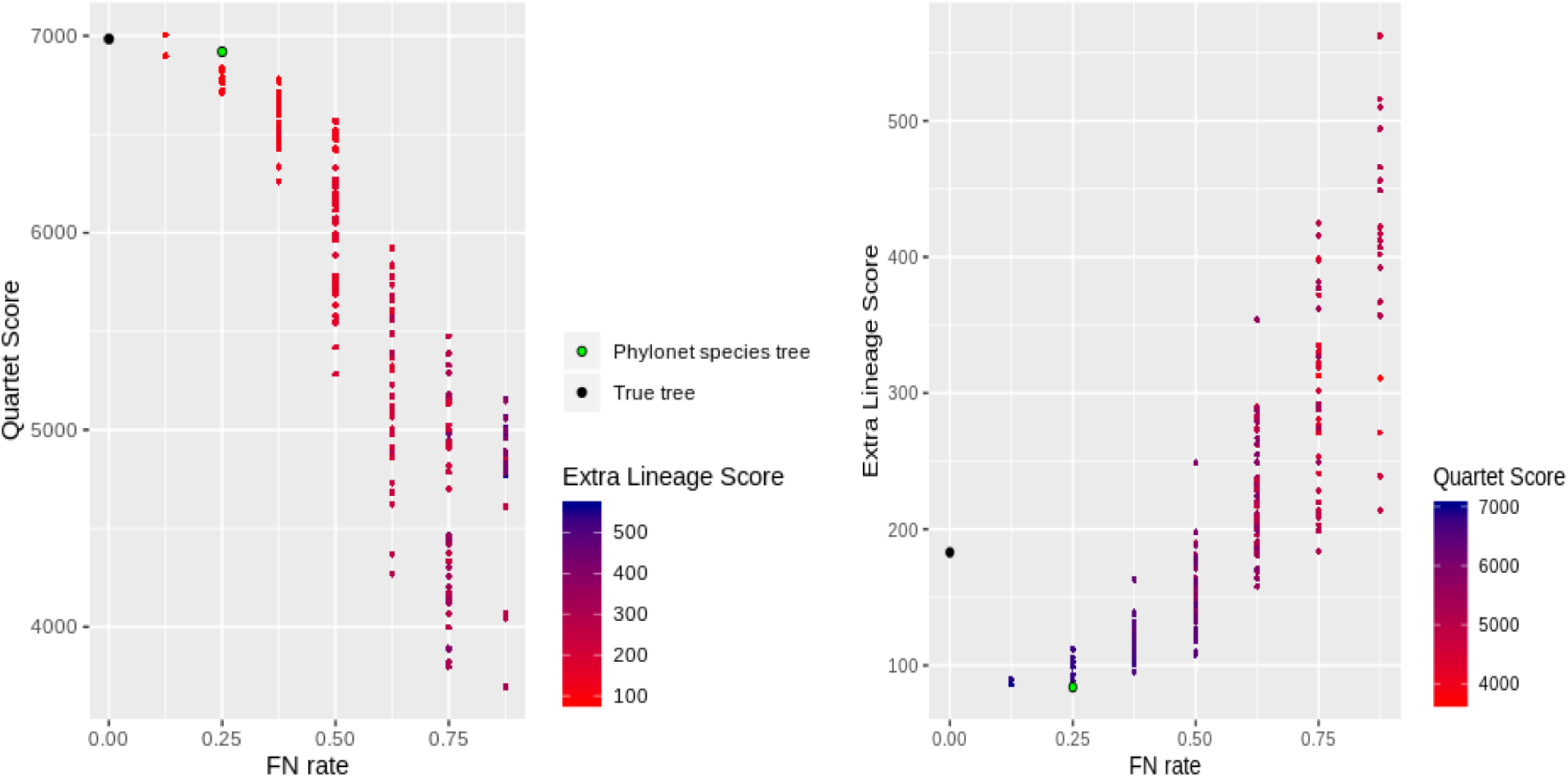
Demonstration of quartet- and MDC-terrace in 11-taxon dataset. Similar to Fig. S9, We show the results for around 250 neighboring trees of the Phylonet-estimated tree.

## Materials and Methods

The following commands were used in this study to run various methods.

### Estimation of species tree by minimizing deep coalescence using Phylonet

The following command was executed in Phylonet to infer species trees from set of gene trees:

~~~
infer st -m MDC -i *<*input gene tree files*>* -x -o *<*output species tree*>*
~~~

The `-x` option was used to specify that the exact version will be run. For the 37-taxon dataset, the default heuristic version was run (without the `-x` option).

### Estimation of species tree by maximizing quartet consistency using ASTRAL-III

The following command was executed in Astral to infer species trees from set of gene trees by maximizing the number of consistent quartets:

~~~
-x -i *<*gene trees*>* -o *<*output species tree*>*
~~~

The `-x` option was used to specify that the exact version will be run. For the 37-taxon dataset, the default heuristic version was run (without the `-x` option).

### Estimation of Extra Lineage Score

The following command was executed in Phylonet to count the total number of extra lineages required to reconcile a set of gene trees in a species tree.

~~~
deep coal count *<*species-tree-file*> <*gene-trees-file*>*
~~~

### Estimation of Quartet Score

The following command was executed in ASTRAL to count the total number of quartets induced by the gene trees that are consistent with a species tree.

~~~
-q *<*species tree to be scored*>* -i *<*gene trees*>* -o *<*output-file*>*
~~~

## Notes

### Competing Interest Statement

The authors have declared no competing interest.

